# Isoform-Dependent Loss- and Gain-of-Function of the Gαs K53N Variant in Human Disease

**DOI:** 10.1101/2025.09.02.673555

**Authors:** Haoran Geng, Justin Martel, Brian Holleran, Andréanne Laniel, Erwan Lanchec, Antoine Désilets, Hugo Giguère, Jean-Luc Parent, Richard Leduc, Louis-Philippe Picard, Mannix Auger-Messier, Christine Lavoie

## Abstract

The K53N mutation in Gαs has been identified in patients with Albright’s Hereditary Osteodystrophy (AHO), pseudohypoparathyroidism type 1A (PHP1a), and dilated cardiomyopathy; however, its molecular mechanism remains unclear. Here, we characterize the molecular, cellular, and physiological consequences of the K53N mutation in both long and short isoform of Gαs. Biochemical analyses reveal that K53N disrupts nucleotide exchange and GTP hydrolysis, rendering both the short (Gαs-S) and long (Gαs-L) isoforms unresponsive to activation by G protein–coupled receptors (GPCRs) or cholera toxin. Both isoforms display a loss-of-function phenotype, failing to trigger cAMP production in response to β2-adrenergic, parathyroid hormone, or vasopressin receptor stimulation. Notably, only the long isoform (Gαs-L K53N) displays constitutive, receptor-independent cAMP generation. The mutation also reduces protein stability, weakens Gβγ subunit interaction, and reduces plasma membrane localization. In neonatal rat ventricular cardiomyocytes, K53N impairs cAMP signaling and exerts dominant-negative effects on isoproterenol-induced responses. Strikingly, only Gαs-L K53N abolishes isoproterenol-stimulated calcium release, directly implicating this isoform in the pathogenesis of cardiomyopathy. Collectively, these findings identify K53N as a unique Gαs mutation that confers both loss- and gain-of-function properties in an isoform-specific manner, providing mechanistic insight into its complex pathogenicity in endocrine and cardiac tissues.

## Introduction

Gαs is a member of the heterotrimeric G proteins family, activated by G protein-coupled receptors (GPCRs) and central in signal transduction processes (*1*). In its inactive state, Gαs is bound to GDP and tightly associated with the Gβγ dimer. Upon agonist stimulation, GPCRs act as guanine nucleotide exchange factors (GEFs), facilitating the exchange of GDP for GTP on Gαs (*2*). This active GTP-bound Gαs dissociates from Gβγ, enabling both subunits to interact with downstream effectors (*3*, *4*). A key effector of Gαs is adenylate cyclase, which converts ATP into the second messenger cAMP, thereby mediating diverse cellular responses (*5*–*7*). Termination of effector stimulation occurs through the intrinsic GTPase activity of Gαs, leading to the reassociation of Gαs-GDP with Gβγ. Ubiquitously expressed, Gαs is encoded by the GNAS gene which comprises 13 exons and 12 introns (*8*). Alternative splicing of exon 3 gives rise to two transcript isoforms, namely the long (Gαs-L) and short (Gαs-S) forms (*8*–*10*). Structurally, both isoforms of Gαs comprises two domains: the alpha helical domain (AHD) and the guanine nucleotide-binding, RAS homology domain (RHD). Activation of Gαs requires separation of these domains, resulting in an “open” conformation that permits GDP release and GTP binding (*11*). Exon 3 encodes a 14-amino acid unstructured segment within a hinge-like region between the AHD and RHD(*12*, *13*). Both isoforms are functional and capable of stimulating adenylate cyclase (*14*, *15*), but a few reports indicate that they differ in their regulatory effects on adenylate cyclase or interaction with distinct effectors, suggesting potential isoform-specific signaling roles (*16*–*18*).

Over 400 genetic mutations within the GNAS locus have been linked to various human diseases (*19*–*21*). Gain-of-function mutations in Gαs have been identified in McCune–Albright syndrome and various cancers (*22*–*24*). Notably, deep sequencing studies have revealed that 4.2% of tumors harbor activating mutations in GNAS (*25*). Heterozygous loss-of-function mutations in Gαs result in Albright’s Hereditary Osteodystrophy (AHO), a syndrome characterized by skeletal and developmental abnormalities such as brachydactyly, short stature, obesity, and mental deficits, among other manifestations (*26*, *27*). An interesting feature of AHO is that maternal inheritance of Gαs loss-of-function mutations leads to multihormone resistance, termed pseudohypoparathyroidism type 1A (PHP1a), while paternal inheritance does not, resulting in pseudopseudohypoparathyroidism (PPHP) (*19*, *28*, *29*). PHP1a is characterized mainly by parathyroid hormone (PTH) resistance leading to decreased serum calcium, increased serum phosphate, and elevated parathyroid hormone (PTH) levels. Some PHP1a patients exhibit resistance to additional hormones acting via other Gαs-coupled GPCRs, including thyroid-stimulating hormone (TSH), growth hormone-releasing hormone (GHRH), and gonadotropins, leading to hypothyroidism, growth hormone deficiency, and hypogonadism (*19*, *21*). The parent-of-origin-specific effect of GNAS mutations arises from the preferential expression of GNAS from the maternal allele in specific endocrine tissues, including the thyroid, gonads, pituitary, and proximal renal tubule (*19*). Since these tissues typically silence the paternally derived allele, a mutation on the paternal allele does not result in hormone resistance in these organs (*19*). As *GNAS* is expressed in a biallelic manner in most tissues, a loss-of-function mutation leads to 50% reduction in Gαs activity, sufficient to maintain a normal signaling activity in most cells, but leading to haploinsufficiency in others, causing AHO features. However, even in tissues with biallelic expression, such as the heart, some studies suggest a preference for the maternal allele (*30*).

To date, over 150 distinct mutations in Gαs have been documented in Albright’s Hereditary Osteodystrophy (AHO), with more than 30% categorized as missense changes (*21*, *31*, https://www.hgmd.cf.ac.uk/ac/all.ph). The molecular basis for the loss of function associated with these mutations includes defective GTP binding, enhanced GTP hydrolysis, impaired GPCR coupling, and decreased protein stability (*21*, *31*–*36*). Recently, a novel GNAS variant, c.159A>G, p.K53N, was identified in a family displaying classical features of AHO and pseudohypoparathyroidism type 1a (PHP-1a) (*37*). Notably, one of the probands with this variant also presented an atypical case of chronic dilated cardiomyopathy, ultimately requiring a heart transplant (*37*). The K53N mutation stands out as unique, as it diverges from any previously documented mutations in Gαs. However, the specific molecular mechanisms driving its pathogenic effects have yet to be explored. Based on the pathological effects observed with this mutant, it was suggested to possess inactivating properties in endocrine tissues and activating properties in cardiac tissues (*37*).

In this study, we investigated the impact of the Gαs K53N variant on the cellular signaling of three Gαs-coupled GPCRs (β2 adrenergic receptor (β2AR), PTH receptor 1 (PTHR1) and vasopressin receptor 2 (V2R)). Our findings reveal that both the short and long isoforms of Gαs K53N are unresponsive to stimulation by these receptors, indicating a clear loss-of-function phenotype. Notably, the long isoform (Gαs-L K53N) displays constitutive, receptor-independent activation of adenylate cyclase, revealing an isoform-specific gain-of-function effect. The mutation also conferred resistance to cholera toxin, weakened Gβγ interaction, reduced protein stability, and altered plasma membrane targeting in both isoforms. In primary neonate rat ventricular cardiomyocytes, expression of either isoform of Gαs K53N results in a markedly diminished cAMP response to isoproterenol and exerts dominant-negative effects on endogenous signaling. Strikingly, only the long isoform of Gαs K53N abolishes isoproterenol-induced calcium release, implicating this variant in cardiomyopathy pathogenesis. Biochemical analyses with purified proteins further reveal that K53N impairs GTP binding and subsequent GTP hydrolysis, providing a mechanistic basis for the observed signaling defects. Together, these findings elucidate how the K53N mutation disrupts Gαs function at molecular, cellular, and physiological levels, offering new insights into its complex pathogenic mechanisms.

## Results

### K53 is a conserved residue located in the P-loop of Gαs Ras-homology domain

Residue K53 (common Gα numbering system corresponding to K53^G.H1.01^ in Gαs-L (*38*)) is found in the β1-α1 loop, also known as the phosphate binding loop (P loop; residues 41-64) of Gαs. Sequence alignment of this region of Gαs reveals that K53 (in yellow) as well as the β1-α1 loop, are very well-conserved amongst Gαs proteins of several species (Fig. 1A). Notably, no sequence differences occur within the 41–64 region between the long (UniProt: P63092-1) and short (UniProt: P63092-2) isoforms. The two isoforms differ only between residues 71–84 (Fig. 1A), a segment in Gαs-L located within a hinge-like region connecting the AHD and RHD (Fig. 1B). This segment is disordered (i.e. unstructured) in available crystal structures (Fig. 1B(i)). The highly conserved β1-α1 loop is part of the Gαs guanosine nucleotide binding pocket and is required for GDP/GTP binding (*39*). Structural modeling reveals that K53 lies in close proximity to residues S54^G.H1.02^, R201^G.hsf2.03^ and D223^G.S3.07^ and can form an electrostatic bond with the γ-phosphate group of the guanine nucleotide (Fig. 1B(ii)), suggesting an important role of K53 in Gαs activity.

**Figure 1:**
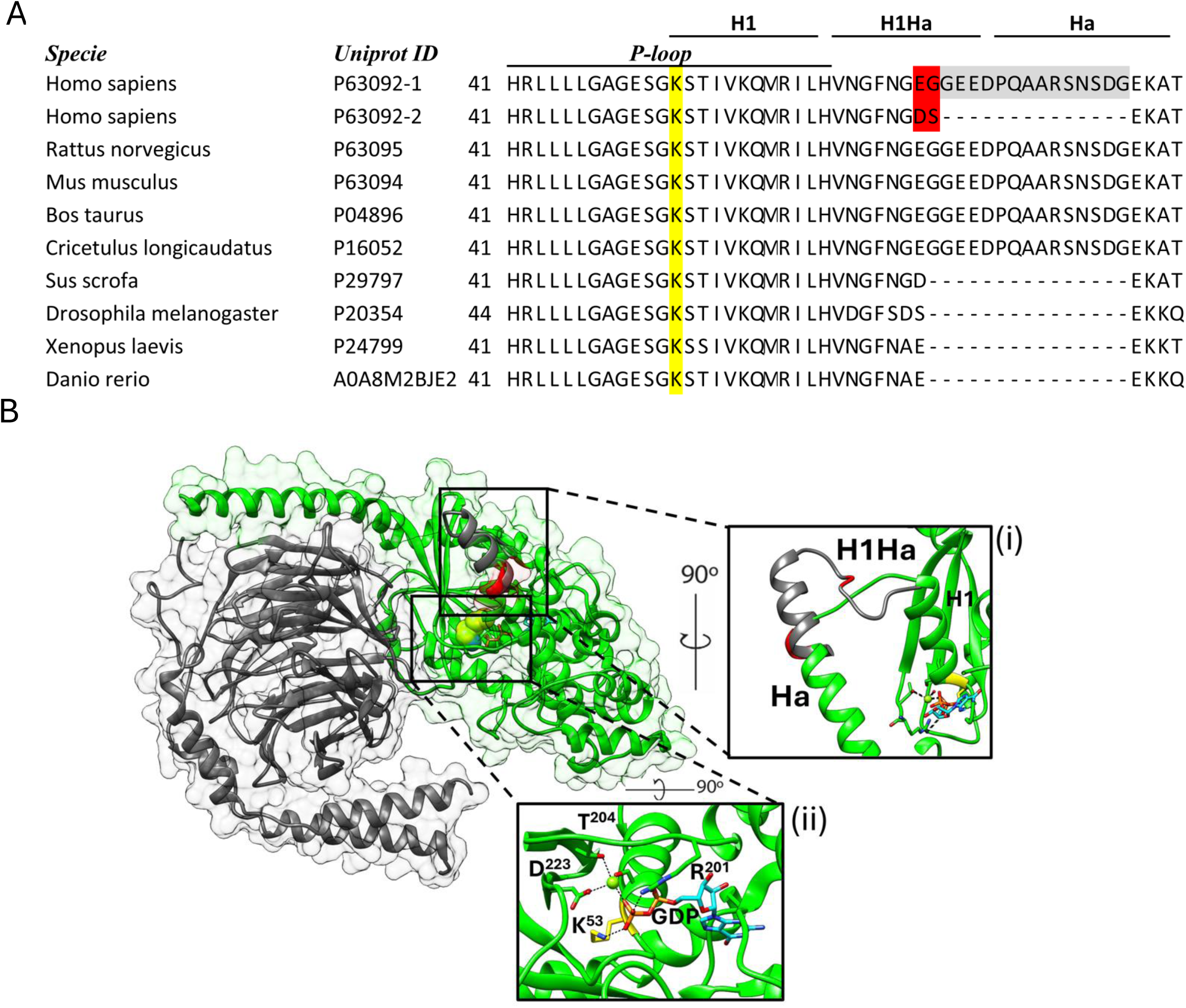
K53 is a highly conserved residue in the P-loop of Gαs Ras-homology domain. (**A**) Sequence alignment of the 41–64 region, along with the additional sequence present in the long (UniProt: P63092-1) and short (UniProt: P63092-2) isoforms of Gαs from Homo sapiens and other species, highlights the high conservation of K53 (yellow). Differences between isoforms in Homo sapiens—two distinct residues (red) and 14 additional residues unique to the long variant (light gray)—are marked. (**B**) Crystal structure of Gαs (green), based on Gαs-S (PDB: 6EG8) and Gαs-L (AF-P63092-F1), displays K53 (yellow) and GDP (orange and blue sticks), with Gβγ subunits shown in gray. Inset (i): The location of the two unique residues (red) and 14 additional residues (gray) in the long isoform, residing within the hinge-like H1Ha segment between the Ras-homology domain (H1) and the α-helical domain (Ha). This grey region is disordered (i.e. unstructured) in crystal structures, with secondary structure predictions (shown) based on AlphaFold. Inset (ii): Structural detail of the nucleotide-binding pocket, highlighting K53 (yellow) forming a hydrogen bond with the phosphate group of GDP (orange).

To characterize the impact of the K53N mutation on Gαs function, we generated K53N variants for both Gαs-S and Gαs-L and transiently expressed them in HEK293 Gαs knock-out cells (HEK Gαs-KO, Fig. 2). Expression levels of Gαs K53N were comparable to WT for both isoforms under steady-state conditions (Fig. 2A), indicating that the mutation does not impair steady state protein stability. However, following cycloheximide treatment to block protein synthesis, both Gαs-S K53N and Gαs-L K53N exhibited accelerated degradation compared to their WT counterparts (Fig. 2B). Densitometric analysis revealed that the half-lives of Gαs WT short and long isoforms were similar at 2.8 and 2.0 hours, respectively (Fig. 2C). However, the half-life of the Gαs-S K53N was reduced to 1.9 hours, and Gαs-L K53N to 0.4 hours. These results demonstrate that substitution of lysine 53 with asparagine, thereby abolishing the positive charge on the sidechain of lysine, markedly destabilizes Gαs proteins— particularly the long isoform—highlighting the critical role of this conserved residue in maintaining protein stability.

**Figure 2.**
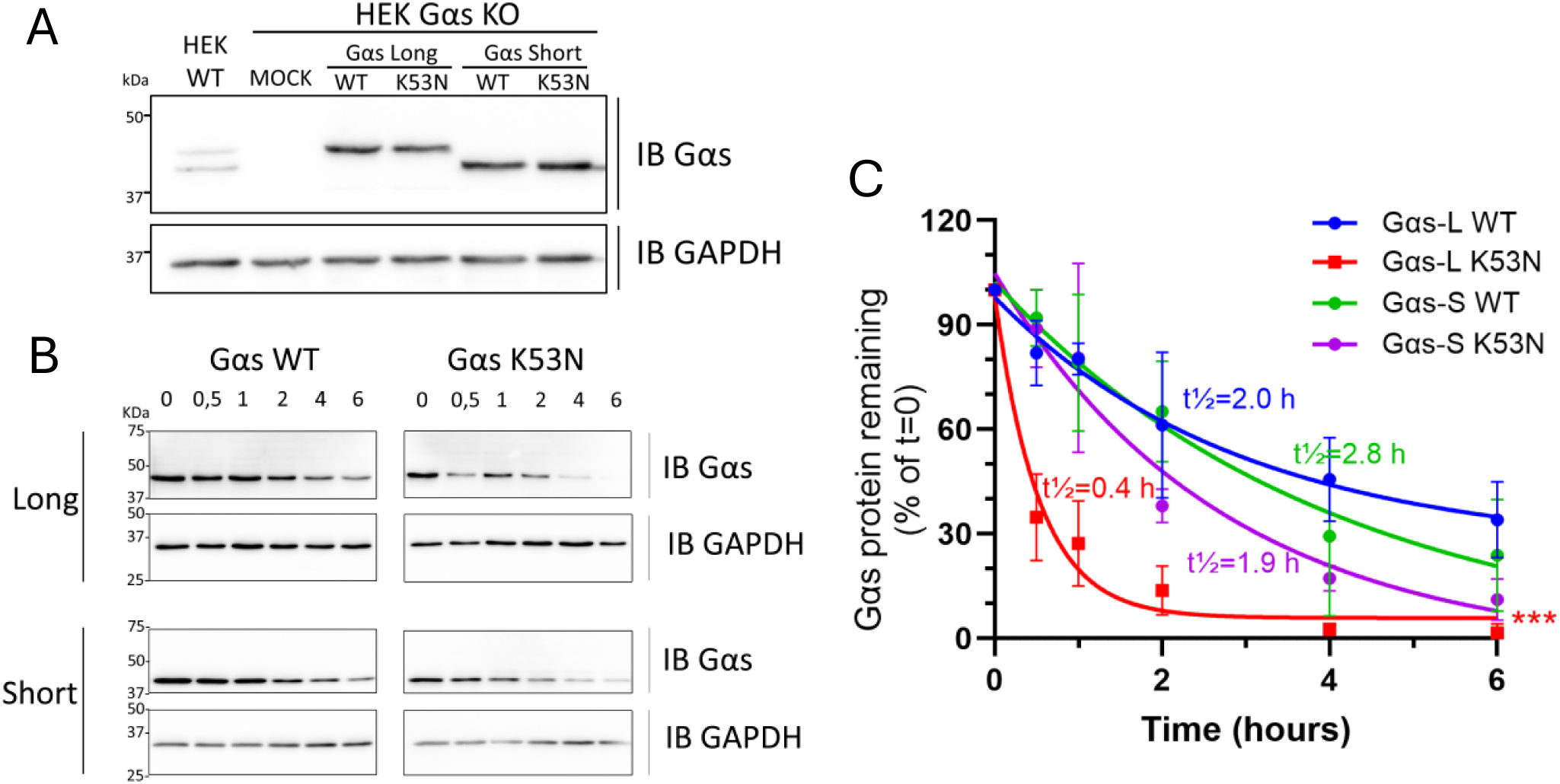
Stability of Gαs K53N compared to Gαs WT. (**A**) Representative Western blot analysis of lysates from HEK Gαs-KO cells transfected with equal amounts of plasmids encoding for Gαs WT or K53N long and short isoforms. GAPDH served as a loading control. Comparison of Gαs WT and Gαs K53N protein half-life. Cells were transfected as in (**A**) and treated with cycloheximide for 0, 0.5, 1, 2, 4 or 6 hours, followed by lysis and western blot analysis. (**C**) Densitometric analysis of panel B. The percentage of remaining Gαs protein at each time point was normalized to the initial (T=0) value and plotted. Data represent mean ± SEM from three independent experiments and were analyzed by Two-way ANOVA followed by Tukey test for multiple comparisons. \**p*<0.03, \**p*<0.002, \*\*\**p*<0.0002, \*\*\*\**p*<0.00001.

### Impact of K53N on cAMP production upon Gαs-coupled GPCR stimulation

The influence of the Gαs K53N mutation was examined on the signaling pathways of three Gαs-coupled G protein-coupled receptors (GPCRs) representing different GPCR classes and physiological relevance. These included the class A prototypical β2 adrenergic receptor (β2AR), given its established expression and function in cardiac tissues (*40*, *41*) and the reported cardiomyopathy in patient expressing Gαs K53N mutation(*37*); the class B parathyroid hormone receptor 1 (PTH1R), whose signaling is altered in AHO and PHP1a (*21*, *42*); and the vasopressin receptor 2 (V2R), another extensively characterized class A GPCR coupled to Gαs whose signaling is unaltered in AHO and PHP1a (*43*–*45*). To assess the ability of Gαs K53N to activate adenylate cyclase and generate cAMP, we employed a bioluminescence resonance energy transfer (BRET)-based assay. This assay utilizes the cAMP-sensing protein ‘effector protein directly activated by cAMP’ (EPAC), which undergoes conformational changes upon cAMP binding, resulting in a reduction of BRET intensity (*46*)(Fig. 3A).

**Figure 3:**
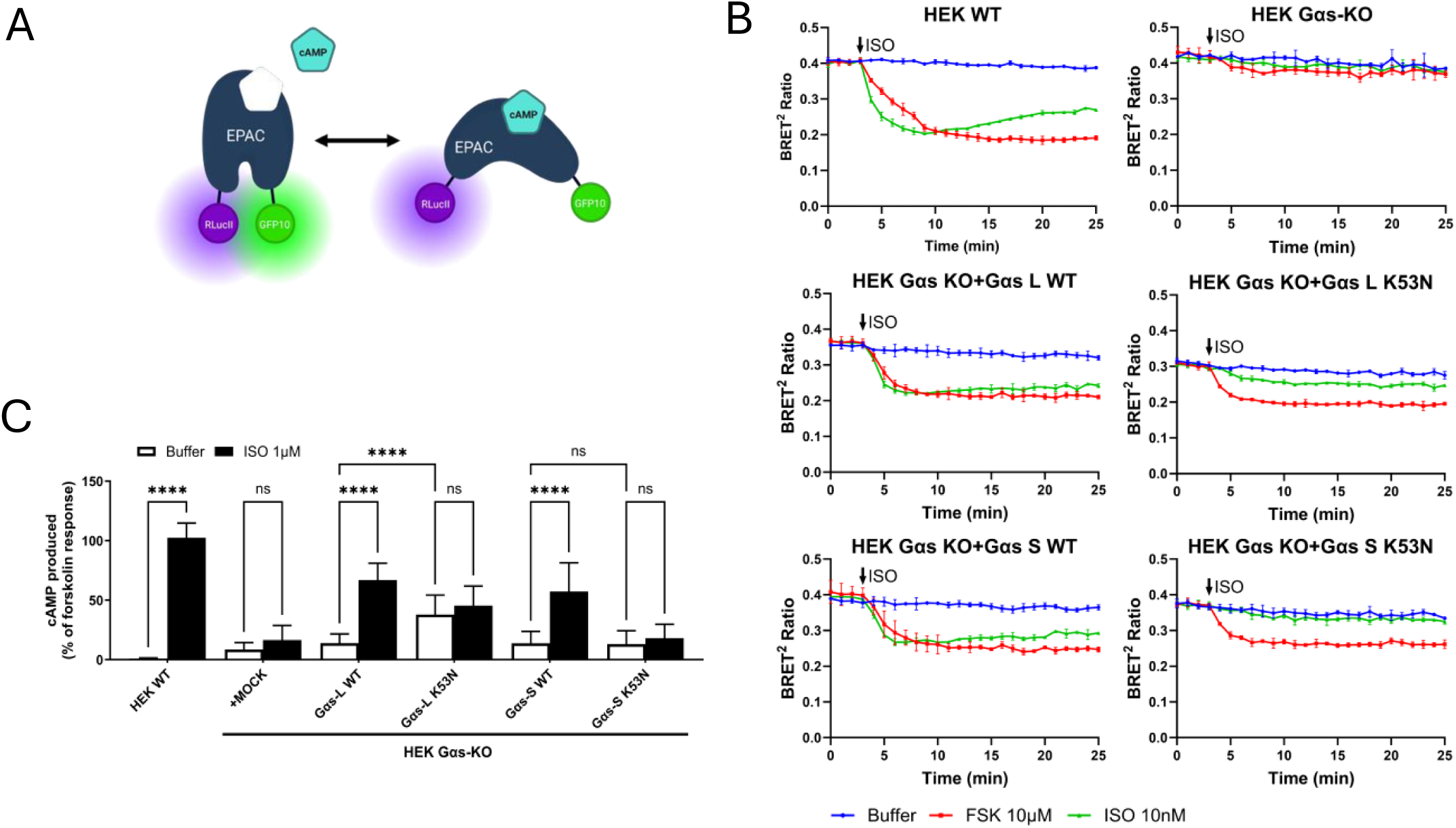
Impact of Gαs K53N mutation on β2AR-induced cAMP generation. (**A**) Schematic representation of the BRET-biosensor EPAC used to detect the intracellular cAMP level. Created with BioRender.com. (**B**) Time-course analysis of cAMP production in HEK293 wild-type (WT) or Gαs-knockout (Gαs-KO) cells transfected with either the long (Gαs-L) or short (Gαs-S) isoform of wild-type (WT) or K53N Gαs. Cells were left untreated (buffer) or stimulated with 10 nM isoproterenol (ISO) or 10 μM forskolin (FSK), and cAMP levels were measured using the BRET assay. All experiments were performed in triplicate and expressed as mean ± SD. (**C**) Quantification of cAMP accumulation after 30 minutes of stimulation, assessed by the HT-FRET cAMP assay. For each condition, cAMP levels were normalized to the response obtained with forskolin. All experiments were performed in triplicate and expressed as mean ± SD. Data were analyzed by one-way ANOVA followed by Tukey test for multiple comparisons. \**p*<0.03, \**p*<0.002, \*\*\**p*<0.0002, \*\*\*\**p*<0.00001, ns=not significant.

In parental HEK cells (HEK WT), stimulation of the endogenously expressed β2AR with isoproterenol (ISO) resulted in a reduction of BRET intensity, consistent with an elevation of cAMP (green line, Fig. 3B). However, in HEK Gαs-KO cells, isoproterenol failed to induce a similar reduction of BRET intensity. Similarly, the adenylyl cyclase activator forskolin did not elevate cAMP levels in HEK Gαs-KO cells (red line, Fig. 3B), supporting previous reports that the affinity of forskolin to adenylate cyclase is greatly reduced in the absence of Gαs, and underscoring the necessity of Gαs for forskolin’s activation of adenylate cyclase (*47*–*50*). When either Gαs-L WT or Gαs-S WT were expressed in HEK Gαs-KO cells, the production of cAMP by adenylate cyclase was restored upon β2AR activation with isoproterenol or forskolin treatment. However, cAMP production profiles were significantly altered with the expression of the K53N mutant. The isoproterenol response was markedly reduced in cells expressing Gαs-L K53N and was completely absent in cells expressing Gαs-S K53N. Additionally, a lower baseline BRET level (blue line) was observed in cells expressing Gαs-L K53N, indicating an elevated basal cAMP level and suggesting higher constitutive activity.

To better characterize and quantify the production of cAMP under basal and agonist-stimulated conditions, we next measured the levels of cAMP using a homogeneous time resolve fluorescence (HTRF) assay in HEK Gαs-KO cells expressing either long or short isoforms of Gαs WT and K53N (Fig. 3C). Isoproterenol stimulated similar accumulation of cAMP in cells expressing Gαs-L WT and Gαs-S WT. However, both isoforms of Gαs K53N were unresponsive to the agonist stimulation, showing levels of cAMP in basal and stimulated condition that were not significantly different. Note that the K53N defect was not associated with reduced expression of the mutant proteins; indeed, immunoblots show that Gαs K53N was expressed at similar level as Gαs WT in these experiments (Fig. S1). This assay also confirmed that the long isoform of Gαs K53N, but not the short one, produces markedly elevated basal levels of cAMP (2.7-fold higher than the basal level of Gαs-L WT). Together, these results suggest that the K53N mutation markedly impairs the ability of Gαs to stimulate cAMP accumulation in response to β2AR stimulation, and that the long isoform of Gαs K53N constitutively activates adenylate cyclase.

To determine whether the effect of Gαs K53N was receptor dependent, we next examined its impact on the signaling of V2R, heterogeneously expressed in cells, using the EPAC biosensor (Fig. 4). In HEK Gαs-KO cells, stimulation of V2R by oxytocin (OT)) did not result in a reduction of BRET intensity compared to HEK293 parental cells (HEK WT) (green line, Fig. 4A). In HEK Gαs-KO cells expressing the long or short isoforms of Gαs WT or K53N, a clear reduction in BRET signal attributable to elevation of cAMP levels can be observed for Gαs WT, but no response was detected with both isoforms of Gαs K53N (Fig. 4A). Quantifying cAMP levels using the HTRF assay (Fig. 4B) further confirmed that both the long and short isoforms of K53N mutant were completely unresponsive to the agonist stimulation, in contrast to Gαs WT. Moreover, the long isoform of Gαs-L K53N mutant resulted in a 3.0-fold increase in basal cAMP levels compared to WT Gαs-L. This effect was not observed with the short isoform Gαs-S K53N. These findings with V2R are similar to those obtained with β2AR.

**Figure 4:**
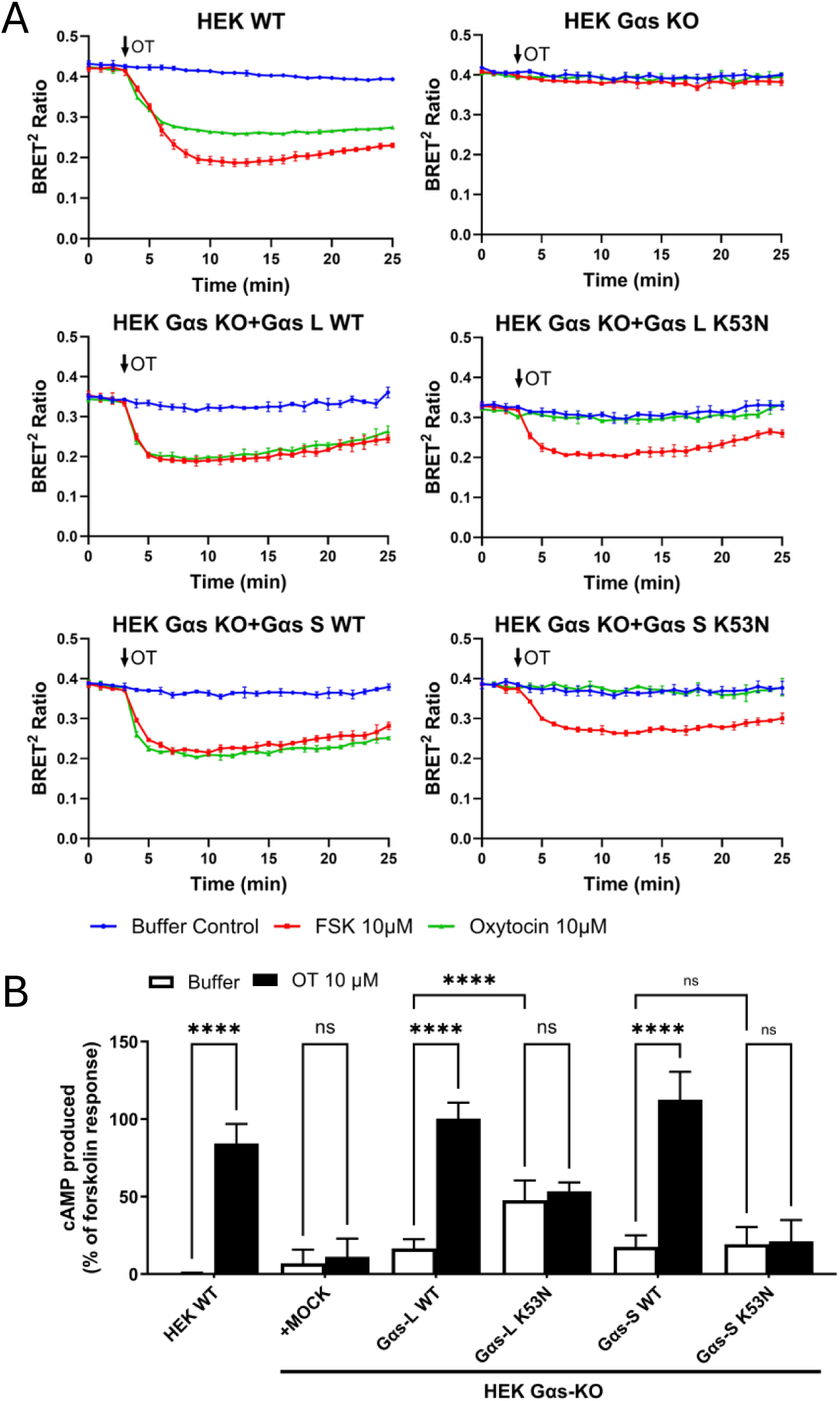
Impact of Gαs K53N mutation on V2R-induced cAMP generation. (**A**) Time-course analysis of cAMP production in HEK293 WT or Gαs-KO cells co-transfected with V2R and either the long (Gαs-L) or short (Gαs-S) isoform of WT or K53N Gαs. Cells were left untreated (buffer) or stimulated with 10 μM oxytocin (OT) or 10 μM forskolin (FSK), and intracellular cAMP levels were monitored using the EPAC-based BRET biosensor assay. All experiments were performed in triplicate and expressed as mean ± SD. (**B**) Quantification of cAMP accumulation after 30 minutes of stimulation, assessed by the HT-FRET cAMP assay. For each condition, cAMP levels were normalized to the response obtained with forskolin. All experiments were performed in triplicate and expressed as mean ± SD. Data were analyzed by one-way ANOVA followed by Tukey test for multiple comparisons. \**p*<0.03, \**p*<0.002, \*\*\**p*<0.0002, \*\*\*\**p*<0.00001, ns=not significant.

Finally, the impact of Gαs K53N mutation was evaluated on PTH1R, a class B GPCR whose impaired signaling is linked to pseudohypoparathyroidism 1A, hypocalcemia, hyperphosphatemia and AHO, conditions observed in patients carrying the K53N mutation in *GNAS* (*37*). PTH1R was heterogeneously expressed in either HEK WT or HEK Gαs-KO cells and cAMP production was measured using EPAC biosensor BRET assay (Fig. 5A) or HTRF assay (Fig. 5B) under both basal and agonist-stimulated conditions. In both assays, cells expressing Gαs-S K53N and Gαs-L K53N failed to respond to stimulation with the PTH1-36 agonist, whereas cells expressing Gαs-S WT and Gαs-L WT produced cAMP in response to the agonist (Figs 5A and B). Additionally, basal cAMP levels were significantly elevated (2.4-fold higher) in cells expressing Gαs-L K53N compared to those expressing Gαs-L WT (Fig. 5B).

**Figure 5:**
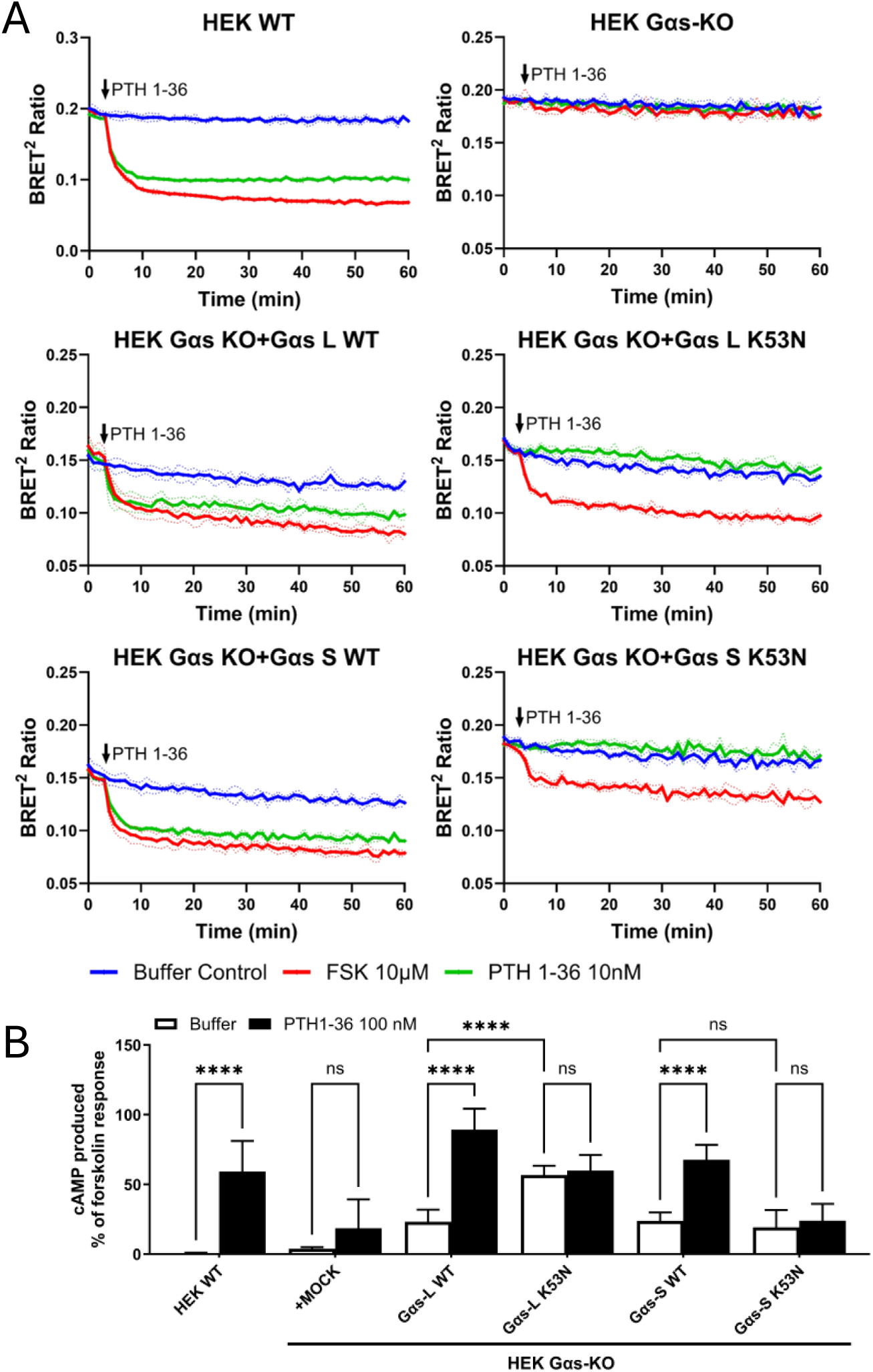
Impact of Gαs K53N mutation on PTH1R-induced cAMP generation. (**A**) Time-course analysis of cAMP production in HEK293 WT or Gαs-KO cells co-transfected with PTH1R and either the long (Gαs-L) or short (Gαs-S) isoform of WT or K53N Gαs. Cells were left untreated (buffer) or stimulated with 10 nM PTH 1-36 or 10 μM forskolin (FSK), and intracellular cAMP levels were monitored using the EPAC-based BRET biosensor assay. All experiments were performed in triplicate and expressed as mean ± SD. (**B**) Quantification of cAMP accumulation after 30 minutes of stimulation, assessed by the HT-FRET cAMP assay. For each condition, cAMP levels were normalized to the response obtained with forskolin. All experiments were performed in triplicate and expressed as mean ± SD. Data were analyzed by one-way ANOVA with Tukey test for multiple comparisons. \**p*<0.03, \**p*<0.002, \*\*\**p*<0.0002, \*\*\*\**p*<0.00001, ns=not significant.

Collectively, these findings indicate that the K53N mutation impairs Gαs activation in response to agonist stimulation across three distinct Gαs-coupled receptors. Notably, the K53N mutation increases basal activity only in the long isoform of Gαs, while the basal activity of the short isoform remains unaffected.

### Effect of K53N on cAMP production under GPCR-independent stimulation of Gαs

Given the unresponsiveness of both long and short isoforms of Gαs K53N to GPCR stimulation, we sought to investigate their direct activation by cholera toxin. Cholera toxin ADP-ribosylates residue R201 of Gαs with slow kinetics, irreversibly inhibiting its GTPase activity and resulting in its constitutive activation independently of GPCR activation (*51*–*53*).

HEK WT and Gαs-KO cells transfected with an empty vector or with the long and short isoforms of Gαs WT or Gαs K53N, were incubated with cholera toxin (CTX), and the kinetics of cAMP production were measured using the EPAC biosensor (Fig. 6). As expected, CTX induced a gradual increase in cAMP levels in HEK WT cells but failed to stimulate cAMP production in Gαs-KO cells. Re-expression of either Gαs-L WT or Gαs-S WT in Gαs-KO cells restored CTX responsiveness, eliciting a similar gradual rise in cAMP levels after ∼1h of stimulation, which continued to increase over the next hour. However, the amplitude and rate of this response were noticeably lower than in HEK WT cells expressing endogenous Gαs. Strikingly, CTX treatment had no effect on cAMP production in Gαs-KO cells expressing either isoform of Gαs K53N. Notably, cells expressing the Gαs-L K53N showed a reduced baseline BRET signal (blue line) compares to Gαs-L WT, consistent with higher constitutive activity. These findings indicate that the inability of Gαs K53N to support both receptor-mediated and CTX-induced cAMP production is unlikely to result from defective GPCR coupling. Instead, the data suggest impaired conformational transitions required for Gαs activation and/or for serving as a substrate for CTX-mediated ADP-ribosylation.

**Figure 6:**
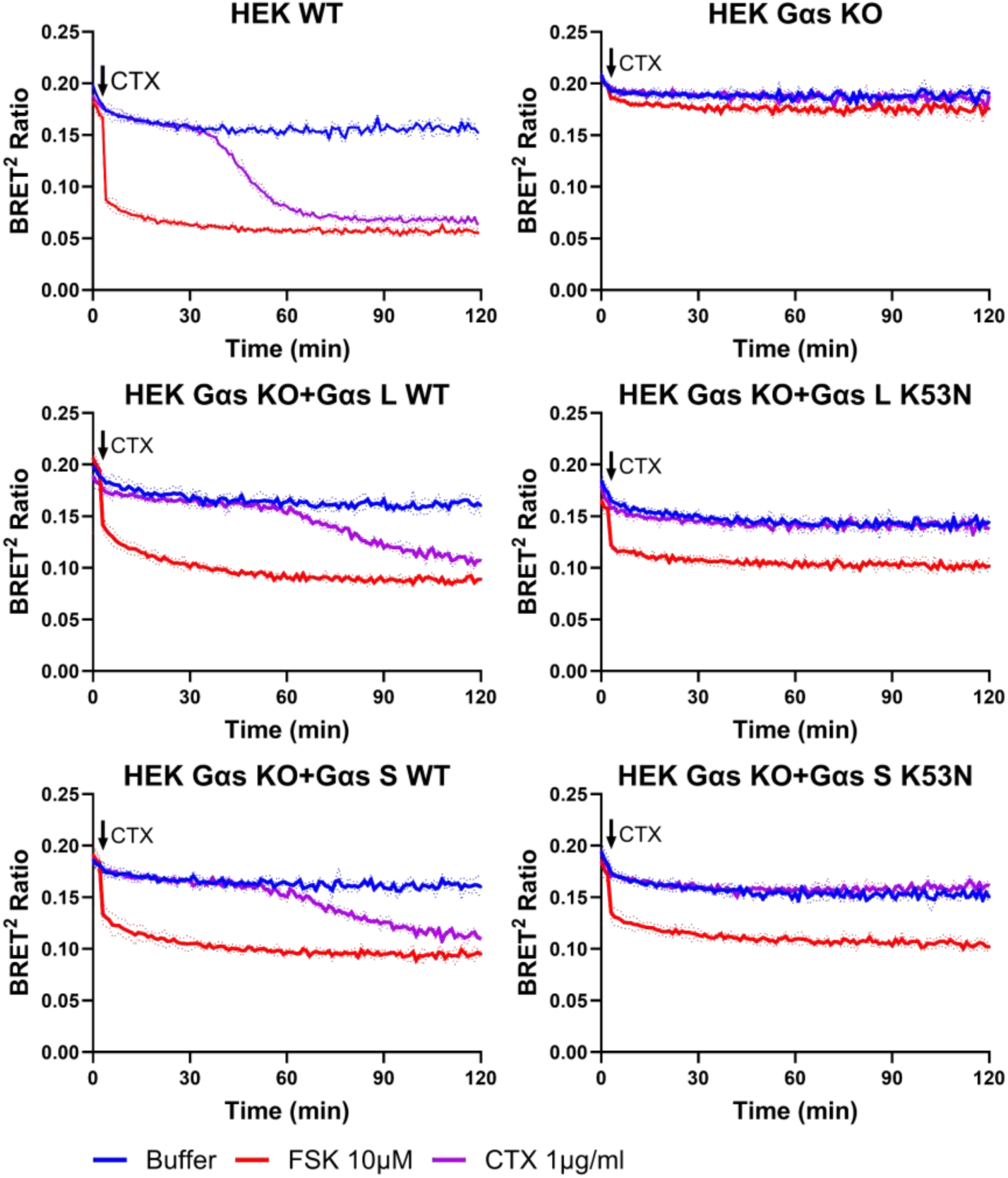
Effect of Gαs K53N mutation on cholera toxin-induced cAMP generation. Time-course analysis of cAMP production in HEK293 WT or Gαs-KO cells transfected with either the long (Gαs-L) or short (Gαs-S) isoform of WT or K53N Gαs. Cells were left untreated (buffer) or treated with 1 ug/mL cholera toxin (CTX) or 10 μM forskolin (FSK), and intracellular cAMP levels were monitored using the EPAC-based BRET biosensor assay. All experiments were performed in triplicate and expressed as mean ± SD.

### Impact of K53N on Gαs interaction with Gβγ and the plasma membrane

We investigated the effect of the K53N mutation on the interaction between Gαs and the Gβγ dimer. A BRET-based biosensor was used to monitor the association of Gαs coupled to renilla luciferase (Gαs-RlucII), with a Gβ1/Gγ1 heterodimer in which Gγ1 was fused to GFP10 (Fig. 7A). In HEK Gαs-KO cells co-expressing β2AR, this biosensor reports both the initial Gαsβγ heterotrimer association and its dissociation upon receptor activation (Fig. 7B). In cells expressing either Gαs-L WT-RlucII or Gαs-S WT-RlucII, baseline BRET confirmed strong heterotrimer association. Isoproterenol stimulation produced a clear decrease in BRET, indicating Gβγ dissociation from activated Gαs WT (Fig. 7B), with statistical analysis confirming a significant reduction at 1□min post-stimulation versus basal condition (Fig. 7C). By contrast, Gαs K53N (both isoforms) displayed a lower baseline BRET signal, consistent with weaker initial heterotrimer assembly, and isoproterenol failed to produce a significant BRET decrease compared to the basal association (Fig. 7B-C). These data suggest that K53N impairs Gαs–Gβγ association, potentially via structural perturbations in the guanine nucleotide-binding pocket.

**Figure 7:**
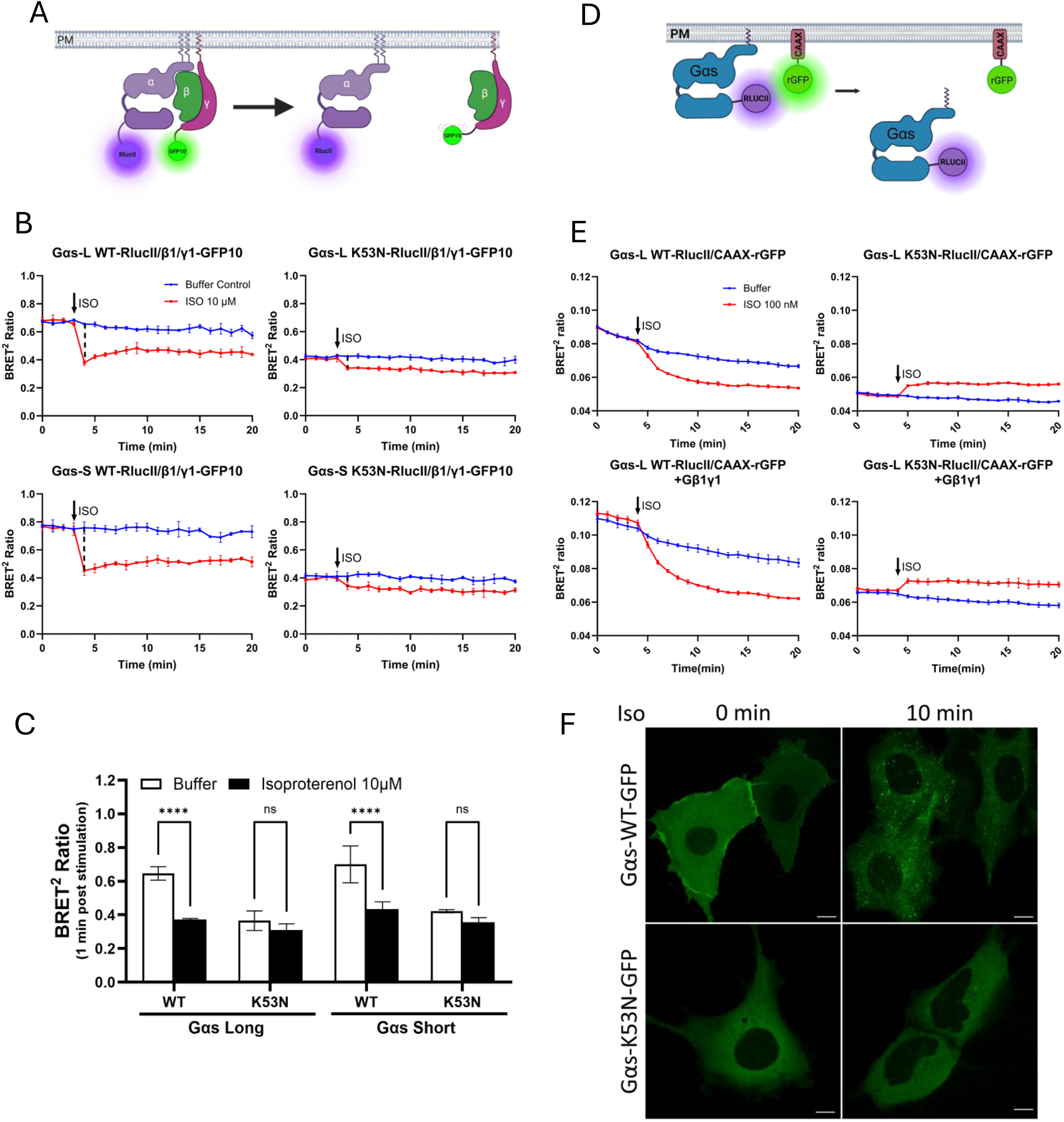
Effect of Gαs K53N mutation on Gβγ interaction and plasma membrane localization. (**A**) Schematic illustration of the BRET-based biosensor used to monitor the association (proximity) of Gαs-RlucII to Gγ-GFP10. Created with BioRender.com. (**B**) HEK Gαs-KO cells were transfected with RlucII-tagged Gαs-L or Gαs-S WT or K53N, along with the Gβ1 and Gγ1-GFP10. Cells were left untreated (Buffer) or stimulated with 10 μM isoproterenol (ISO). Data shown are representative of three independent experiments, each performed in triplicate, and are presented as mean ± SD. (**C**) BRET2 ratios analysis from panel B, comparing Gαs WT and K53N (long and short isoforms) after 1 minute of buffer or ISO treatment. Data were analyzed by one-way ANOVA with Tukey test for multiple comparisons. \**p*<0.03, \**p*<0.002, \*\*\**p*<0.0002, \*\*\*\**p*<0.00001, ns=not significant. **(D)** Schematic of Gαs-RlucII/CAAX-rGFP biosensor. Created with BioRender.com. **(E)** HEK293 cells were transfected with Gαs-L WT-RlucII or Gαs-L K53N-RlucII together with CAAX-rGFP. After 48h, cells were left untreated (Buffer) or stimulated with 100 nM of isoproterenol (ISO). Data shown are representative of three independent experiments, each performed in triplicate, and are presented as mean ± SD. **(F)** HEK293 cells were transfected with Gαs-WT-GFP or Gαs-K53N-GFP together with Flag-β2AR and stimulated with 10 μM of ISO for 0 or 10 min. Cells were analyzed by live confocal microscopy. Scale bars, 10 μm.

We next examined whether the K53N mutation alters Gαs localization at the plasma membrane (PM). HEK293 cells expressing β2AR were transfected with Gαs-RlucII and a PM-anchored rGFP containing a prenylated CAAX motif (rGFP–CAAX) to monitor membrane association via ebBRET (Fig. 7D) (*54*). In cells expressing Gαs-L WT, β2AR activation by isoproterenol induced a rapid decrease in BRET signal (Fig. 7E), consistent with dissociation from the PM and in line with previous reports (*55*, *56*). In contrast, Gαs-L K53N displayed a markedly lower baseline BRET signal, indicative of weaker initial membrane association. Strikingly, isoproterenol stimulation led to an increase—rather than a decrease—in BRET signal, suggesting recruitment of the mutant protein to the PM upon receptor activation. Co-expression of Gβ1/Gγ1 enhanced both the basal PM association of Gαs-L WT and its agonist-induced dissociation. For Gαs-L K53N, Gβ1/Gγ1 co-expression slightly increased basal PM association but did not significantly alter its recruitment profile following receptor stimulation, supporting earlier findings that K53N disrupts Gαs-Gβγ interactions.

Live cell confocal microscopy was employed to validate the cellular localization of Gαs K53N, tagged with GFP. GFP was strategically inserted into the loop encoded by exon 3 of Gαs-L, replacing the amino acids 73-84. This insertion is known to maintain the functional activity of Gαs (*57*). Consistent with previous reports, wild-type Gαs-GFP was observed in the cytoplasm, with a predominant presence at the PM (Fig. 7F). Upon stimulation with isoproterenol, a notable redistribution of Gαs-GFP to intracellular vesicles was observed. In contrast, K53N Gαs-GFP exhibited a predominantly cytoplasmic localization, with minimal presence at the PM, and this distribution remained unaltered following agonist stimulation. These findings underscore the significant impact of the K53N mutation on Gαs intracellular localization and its correlation with the diminished association with Gβγ and the PM.

Together, these findings indicate that the K53N mutation severely impairs Gαs–Gβγ assembly, plasma membrane association, and receptor-driven trafficking, most likely through mislocalization and/or weakened interaction, potentially influenced by altered nucleotide state or protein conformation.

### Validation of the impact of K53N mutation in rat primary cardiomyocytes

Given our results and the reported association of the Gαs K53N mutation with cardiomyopathy in a reported patient(*37*), we utilized isolated neonatal rat ventricular myocytes (RNVM) as a model to investigate the cardiac implications of the Gαs K53N mutation. Adenovirus-mediated delivery of short hairpin RNA (shRNA) was used to initially deplete endogenous Gαs. Subsequently, Gαs WT or the K53N mutant was reintroduced into these cardiomyocytes, alongside the EPAC biosensor, to assess the mutation’s impact on Gαs function in these cells (Fig. 8). Western blotting confirmed that efficient depletion of Gαs upon infection with the shGαs adenovirus (Fig. 8A). Upon reintroduction, both the long and short forms of Gαs WT or K53N were detected at the expected molecular weights. The variation in expression levels of the K53N mutant likely reflects the mutation’s detrimental impact on Gαs protein stability, consistent with earlier findings presented in Fig. 2. Monitoring cAMP levels via the EPAC biosensor revealed a significantly reduced, though not completely abolished, cAMP response to isoproterenol stimulation in shRNA Gαs-infected cells compared to shRNA control cells (Fig. 8B, C). Reintroduction of either Gαs WT isoform fully restored cAMP production, whereas cells expressing Gαs K53N exhibited a markedly reduced response compared with Gαs WT. Notably, the cAMP levels in cells expressing Gαs K53N were significantly lower than those in shRNA Gαs-treated cells (Fig. 8C), suggesting a dominant negative effect of the mutant.

**Figure 8:**
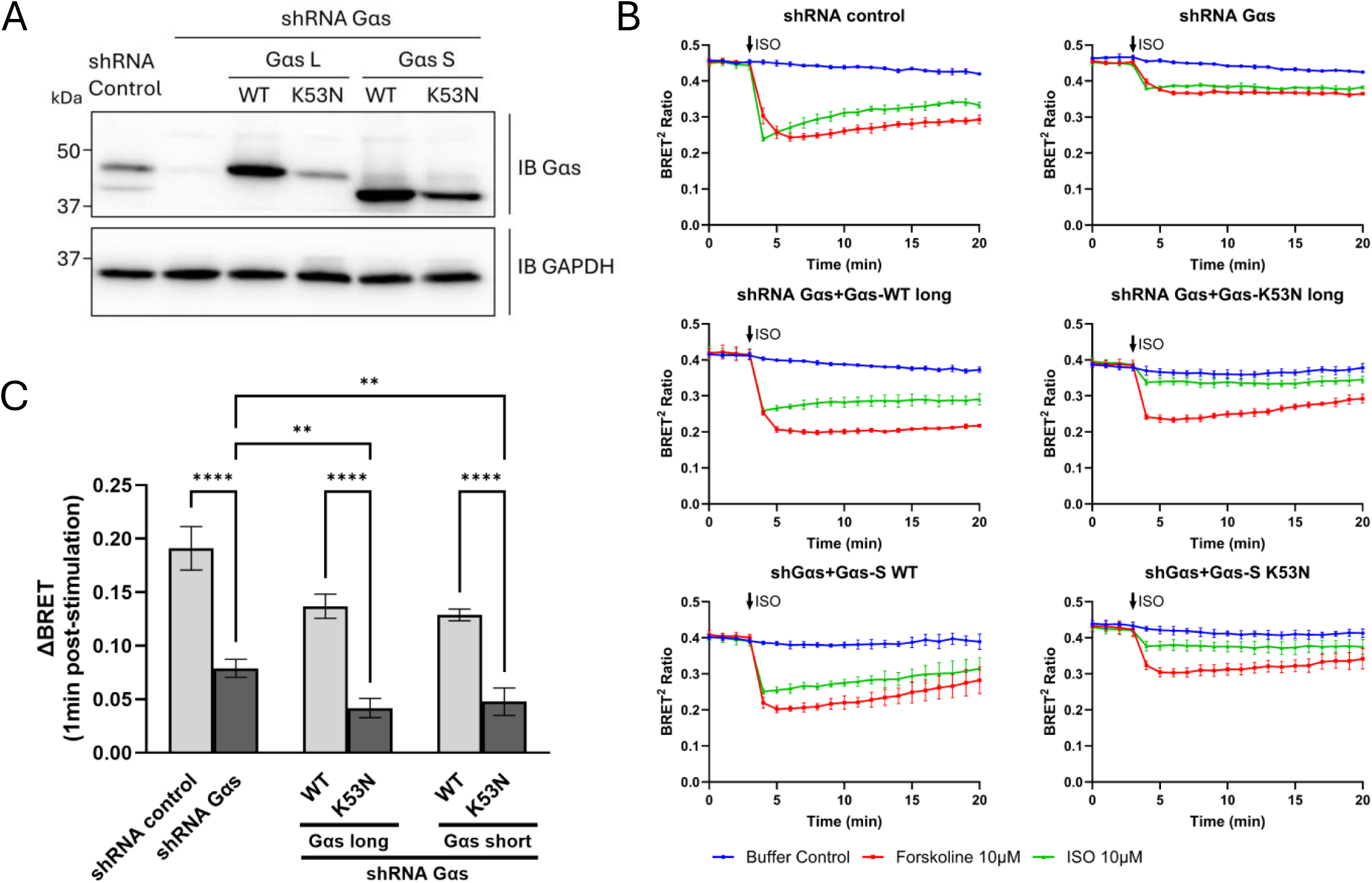
Impact of Gαs K53N on cAMP production in primary neonatal rat ventricular myocytes. (**A**) Representative Western blot analysis of lysates from NRVM. Cells were infected with either control shRNA or shRNA targeting Gαs, followed by re-infection with Gαs WT or K53N (long or short isoform). Lysates were immunoblotted with anti-Gαs or anti-GAPDH antibodies. (B) Time-course analysis of cAMP production in NRVM treated as in (A) and co-expressing the EPAC biosensor. Cells were either untreated (Buffer) or stimulated with 10 μM isoproterenol (ISO) or 10 μM forskolin (FSK), and cAMP levels were measured using a BRET assay. Data represent the mean ± SD from three independent experiments, each performed in triplicate. (**C**) Quantification of BRET2 ratios from panel (**B**) at 1-minute post-treatment with buffer or ISO. Data were analyzed by one-way ANOVA with Tukey test for multiple comparisons. \**p*<0.03, \**p*<0.002, \*\*\**p*<0.0002, \*\*\*\**p*<0.00001, ns=not significant.

We next investigated the ligand stimulated calcium (Ca^2+^) mobilization of NRVM, the process linking electrical excitation at the cell membrane to the production of cytosolic Ca^2+^ signals that trigger cell contraction (*58*, *59*). NRVM were infected with adenoviral vectors carrying control or Gαs shRNA, alongside constructs for the short or long isoforms of Gαs WT or Gαs K53N, or β-galactosidase (as a control). Using fura-2-based calcium imaging with electrical pacing, we measured baseline calcium transient amplitudes (blue line, Fig. 9A) followed by isoproterenol-stimulated transient amplitude (ISO, red line) in individual cells under each condition. Baseline curves were consistent across all groups. However, isoproterenol treatment led to a consistent increase in Ca^2+^ release in all cells except those expressing Gαs-L K53N mutant, which exhibited a notably reduced response. Fig. 9B quantifies the change in transient area under the curve before and after isoproterenol-treatment (as shown in Fig. 9A), averaged across 20 cells from 5 independent experiments. Isoproterenol response was unchanged in Gαs-depleted cells and cells expressing Gαs-L WT, Gαs-S WT, or Gαs-S K53N. In contrast, the Gαs-L K53N mutant significantly impaired the isoproterenol-induced increase in Ca^2+^ transients, suggesting that this mutant alters the calcium mobilization in NRVM.

**Figure 9:**
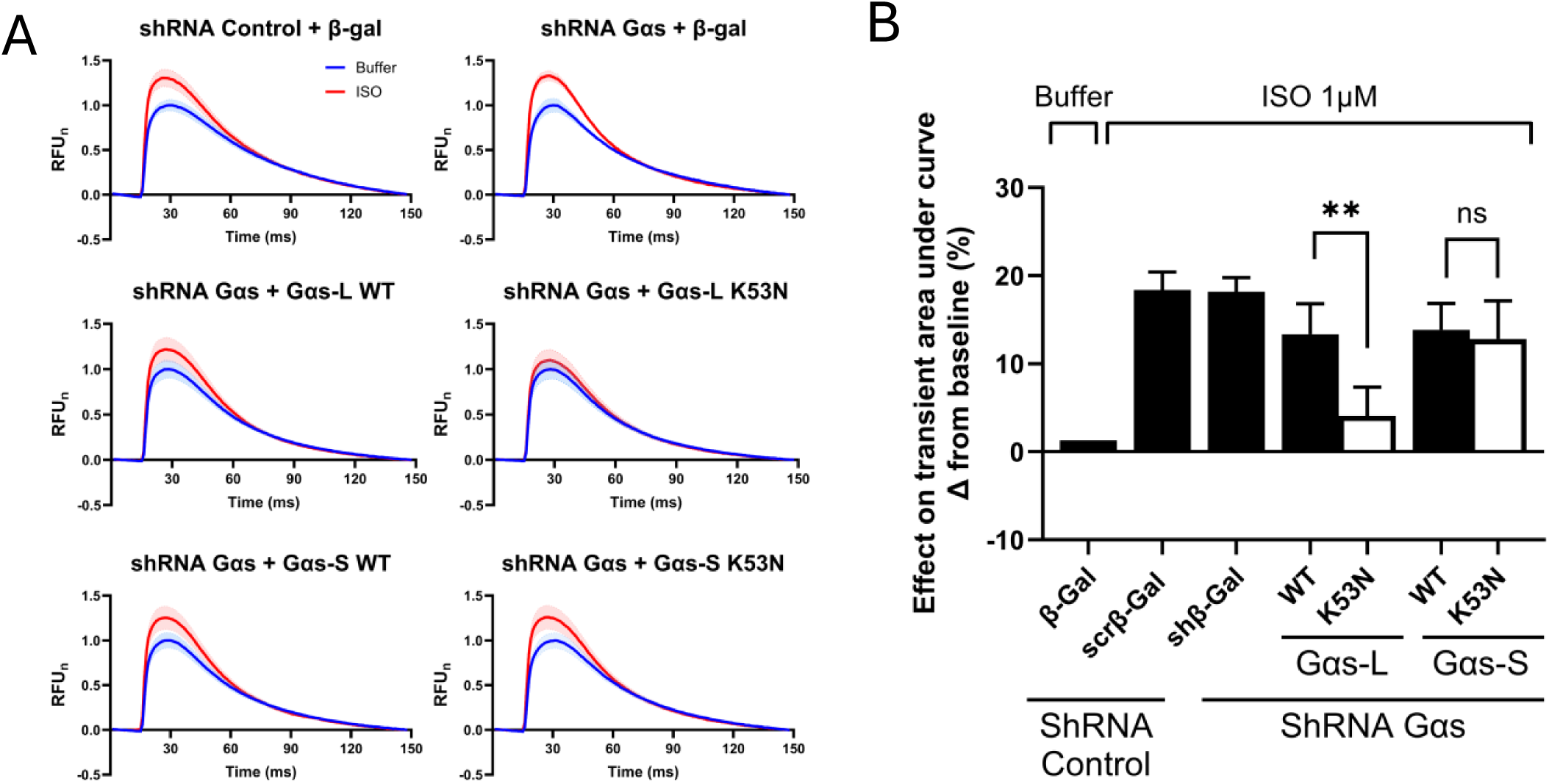
Effect of Gαs K53N mutation on isoproterenol-induced Ca^2+^ release in primary neonatal rat ventricular myocytes. (**A**) NRVMs were infected with either control shRNA or shRNA targeting Gαs, then re-infected with β-galactosidase (β-gal, control), or with WT or K53N mutant forms of the long (Gαs-L) or short (Gαs-S) Gαs isoforms. Cells were loaded with the calcium indicator Fura-2 AM. Shown are representative normalized fluorescence traces (RFUn) from single NRVMs paced at 1.7 Hz, recorded at baseline (blue) and following isoproterenol (ISO) stimulation (red). (**B**) Quantification of the area under the curve (AUC) for Ca²⁺ amplitude traces as shown in (A), from 20 cells per group pooled from 5 independent experiments. Data are presented as the change from baseline. Statistical analysis was performed using one-way ANOVA with Tukey test for multiple comparisons, \**p*<0.03, \**p*<0.002, \*\*\**p*<0.0002, \*\*\*\**p*<0.00001, ns=not significant.

Together, these results show that Gαs K53N acts as a dominant-negative in cardiomyocytes, blunting β-adrenergic cAMP signaling and—specifically in the long isoform—impairing Ca^2+^ transients, providing a plausible link to its association with cardiomyopathy.

### Impact of K53N mutation on GTP binding and hydrolysis

To investigate the biochemical impact of the K53N mutation, we expressed and purified the short isoform of Gαs WT and K53N proteins for enzymatic analysis, as the long isoform proved unstable and could not be purified for *in vitro* characterization (Fig. 10). We first assessed nucleotide exchange by monitoring BODIPY-GTPγS fluorescence. Upon Gαs-S WT addition, BODIPY-GTPγS exhibited rapid fluorescence increases, indicating efficient GTP binding, whereas Gαs-S K53N showed markedly slower binding, reaching only ∼50% of Gαs WT fluorescence after 20 minutes (Fig. 10A), demonstrating impaired GDP/GTP exchange. We next measured intrinsic GTPase activity using a bioluminescent assay (Fig. 10B). Gαs-S K53N displayed reduced cumulative GTP hydrolysis versus Gαs-S WT at equivalent protein concentrations, but with similar kinetic parameters (t½=23 min for Gαs WT and 26 min for Gαs K53N), indicating that GTP hydrolysis itself was not substantially affected. Since this assay reflects both nucleotide exchange and hydrolysis rates, the diminished steady-state GTPase activity of Gαs-S K53N primarily stems from defective nucleotide exchange rather than impaired catalysis.

**Figure 10:**
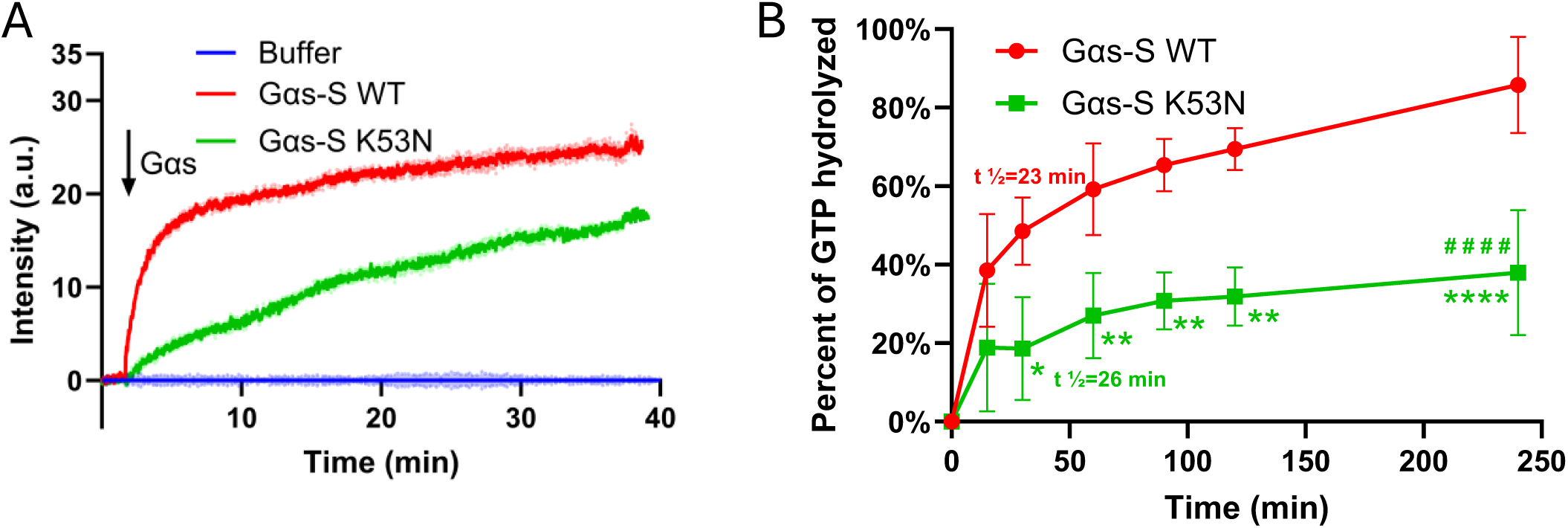
Effect of Gαs K53N mutation on nucleotide exchange and GTP hydrolysis. (A) Nucleotide exchange rate was measured with BODIPY FL GTPγS. Baseline fluorescence (buffer) was established with 100 nM BODIPY FL GTPγS. After a 100-second stabilization, 200 nM of either Gαs short WT (Gαs-S WT) or K53N (Gαs-S K53N) was added, and fluorescence intensity was continuously monitored. Data represent mean ± SD from four independent experiments. (**B**) Time course of GTP hydrolysis by purified Gαs-S WT and Gαs-S K53N was measured using the GTPase-Glo assay at 15, 30, 60, 90, 120, and 240 minutes. Reactions contained 1.25 μM rGTP and 1 μM Gαs protein; a no-protein control defined 0% GTP hydrolyzed. Data are shown as mean ± SD from three independent experiments. Statistical analysis was performed using two-way ANOVA. Significance is indicated as follows: \**p*<0.03, \*\**p*<0.002, \*\*\**p*<0.0002, \*\*\*\**/####p<*0.00001. (*) denote differences between K53N and WT at individual time point; (#) indicate overall differences between the K53N and WT curves.

## Discussion

In this study, we uncover the unique and multifaceted pathogenic mechanisms of the newly identified Gαs K53N variant, found in patients with Albright’s Hereditary Osteodystrophy (AHO), pseudohypoparathyroidism type 1A (PHP1a), and notably, dilated cardiomyopathy (*37*)—a novel clinical association for Gαs mutations. Our results reveal that the single K53N substitution disrupts nucleotide exchange, resulting in both gain- and loss-of-function effects, alongside dominant-negative activity. Remarkably, these outcomes are isoform-dependent: although both Gαs K53N isoforms fail to respond to GPCR stimulation, only the long isoform (Gαs-L K53N) displays paradoxical constitutive activity, enhancing ligand-independent signaling at PTH1R, β2AR, and V2R. Furthermore, only the long isoform Gαs-K53N perturbs calcium signaling in cardiomyocytes, directly implicating this isoform in cardiomyopathy pathogenesis. This study classifies K53N as a unique loss-of-function mutant exhibiting isoform-dependent dual functionality in signaling pathway and pathogenesis.

At the protein structural level, K53 resides within the P-loop of Gαs’ guanine nucleotide-binding pocket—a highly conserved residue amongst Gαs proteins of several species and across all G protein subtypes (Fig. S2). The positive charge of lysine 53 stabilizes critical salt bridges and electrostatic interactions with the β- and γ-phosphates of GDP/GTP (Fig 1B(ii)). Substitution with asparagine (K53N) disrupts these interactions, leading to impaired nucleotide coordination. This is reflected by the markedly slower nucleotide exchange of purified Gαs-S K53N, suggesting reduced affinity for GTP. Correspondingly, the mutant displays reduced GTPase activity, with total hydrolyzed GTP reaching saturation at only 38% for Gαs-S K53N versus 85% for Gαs-S WT, consistent with impaired nucleotide exchange, and suggesting greater loss of affinity for GTP relative to GDP. Nevertheless, the comparable hydrolysis half-time between mutant and WT proteins shows that the intrinsic GTPase activity of Gαs-S K53N remains intact. Notably, a homologous lysine-to-asparagine mutation in Gαo (K46N)—a loss-of-function variant linked to severe neurological disease (*60*)— similarly impairs both GTP binding and hydrolysis (*61*), reinforcing the critical role of the P-loop lysine and its positive charge in nucleotide exchange.

At the cellular level, both the short and long isoforms Gαs K53N fail to support cAMP production in response to stimulation of three prototypical Gαs-coupled GPCRs (β2AR, PTH1R, V2R), when expressed in HEK Gαs-KO cells. This loss-of-function phenotype is paralleled by incapacity to be activated by cholera toxin, reduced association with Gβγ and disrupted plasma membrane localization, further diminishing the ability to form functional heterotrimers and propagate downstream signaling. These findings are consistent with the biochemical studies indicating the nucleotide exchange defect locks Gαs in an inactive or nucleotide-free state, precluding activation by GPCRs or cholera toxin and leading to reduced protein stability.

An intriguing finding is the isoform-specific gain-of-function observed with the Gαs-L K53N mutant, which shows elevated basal cAMP production independent of GPCR activation. Notably, this constitutive activity is absent in the short isoform, and Gαs-L K53N remains unresponsive to agonist stimulation, highlighting that the mutation’s gain-of-function does not rescue normal signaling. While both Gαs splice variants are generally considered functionally equivalent(*16*), prior studies have reported that the long isoform has a higher GDP dissociation rate and marginally enhanced cellular activity, but similar GTP hydrolysis rates (*18*, *62*–*64*). In our cellular assays, however, WT Gαs-L and Gαs-S exhibited comparable activity, suggesting that the K53N mutation uniquely confers this isoform-specific phenotype. There is limited research directly comparing the impact of disease-associated mutations on both Gαs isoforms, and only a handful of mutations have been shown to induce isoform-specific effects in human pathology. Recent RNA-omics studies have highlighted that increased expression of Gαs-L, driven by alternative splicing of GNAS, can promote aberrant signaling and pathological phenotypes in neoplasms and hematopoietic disorders (*18*). Additionally, the well-characterized oncogenic R201 mutation, a classic gain-of-function variant, has been shown to significantly enhance the activity of the long isoform, suggesting a nonredundant and potentially synergistic effect between mutation and isoform context (*18*). Our findings add a new dimension to this paradigm by demonstrating that the loss-of-function K53N mutation paradoxically increases basal activity only in the long isoform. This is reminiscent of the loss-of-function mutations F376V and S54N, both unable to mediate hormone-dependent activation, and notable for increasing ligand-independent activity of Gαs-L, although these studies did not directly compare the effects on Gαs-L versus Gαs-S (*65*, *66*). The F376V mutation, located in the α5 helix, is thought to induce conformational changes that mimic receptor activation, resulting in multisystem disorders such as nephrogenic syndrome of inappropriate antidiuresis and precocious puberty (*65*). In contrast, the S54N mutation, situated in the P-loop adjacent to K53, increases the nucleotide exchange rate (*66*). This variant was engineered based on homology to Ras-S17N, which is known to impair GTP binding in Ras (*67*), but S54N itself is not associated with any pathologies. Other classic pathological loss-of-function mutations, such as R228C in the nucleotide-binding pocket, have not been reported to cause constitutive activation or to display isoform-dependent effects. Further studies comparing the impact of various loss-of-function mutations on the short and long Gαs isoforms would be valuable for understanding isoform-specific functionality.

When expressed in neonatal rat ventricular cardiomyocytes (NRVMs), both short and long isoforms of Gαs K53N failed to restore cAMP signaling during β-adrenergic stimulation and actively suppressed endogenous Gαs activity, demonstrating a dominant-negative effect. Notably, similar dominant-negative properties have been observed for Gαs S54N and Gαo K46N mutants, indicating a conserved mechanism among P-loop mutations (*61*, *66*). This dominant suppression is generally attributed to the mutant protein’s ability to sequester key signaling partners—such as GPCRs, Gβγ subunits, or effectors—thereby preventing activation of Gαs WT, as established for other dominant-negative G protein variants (*68*). Specifically, the K53N mutation reduces Gαs-Gβγ association both under basal conditions and following agonist stimulation, does not interfere with forskolin-induced adenylyl cyclase activation, and biochemically traps Gαs in a nucleotide-free state. These findings suggest that the dominant-negative action of K53N primarily involves sequestration of activated GPCRs by nucleotide-free Gαs, effectively blocking receptor-mediated signaling. The enhanced recruitment of Gαs K53N to the plasma membrane following β2AR activation, as detected by the CAAX-BRET assay, is consistent with this model. Pathologically, the dominant-negative behavior of Gαs K53N would result in severe inhibition of receptor-driven pathways—even in the presence of a functional wild-type allele—ultimately leading to profound disruption of cellular signaling and a disease phenotype with distinctive features.

Interestingly, while both short and long isoforms of Gαs K53N abolish cAMP production in NRVMs, only the long isoform eliminates the isoproterenol-induced increase in cytosolic calcium at the level of excitation–contraction coupling. This finding points to a unique pathological role for Gαs-L in cardiomyocytes. One possible explanation lies in the compartmentalization of cAMP signaling, as PKA activity is highly localized, anchored to specific subcellular sites, and limited to targets within a 15–25 nm radius, allowing for distinct physiological outcomes even at similar global cAMP levels (*66*, *69*–*72*). In cardiomyocytes, such local cAMP and PKA pools are crucial for precise regulation of calcium handling at the L-type Ca^2+^ channel on the PM and ryanodine receptor and SERCA on the sarcoplasmic reticulum and Golgi apparatus (*69*, *73*), and disruption of this spatial organization contributes to cardiac dysfunction (*74*, *75*). The Gαs-L isoform may possess unique structural features or trafficking properties that enable it to scaffold the signaling complexes necessary for β-adrenergic regulation of calcium mobilization—a function not compensated by the short isoform. Supporting this, Gαs isoforms have been reported to redistribute differently between the plasma membrane and light-density membrane fraction or cytosol during β-adrenergic stimulation in both s49 lymphoma cells (*16*, *63*) and rat cardiac tissue (*76*), highlighting the need to further investigate isoform-specific localization and interactions in cardiomyocytes. Another explanation is that Gαs-L might influence calcium handling through cAMP/PKA-independent pathways; for instance, Gαs-L, but not Gαs-S, has been reported to promote β-adrenergic receptor-mediated ERK activation from endosomes (*17*, *18*). Interestingly, ERK activation has been reported to upregulate calcium channel Ca_V_α2δ1 following chronic β1AR stimulation in NRVMs (*77*). Elucidating these mechanisms will be the focus of future investigations.

The loss-of-function and dominant-negative effects of K53N mutation on both Gαs isoforms, directly account for the clinical manifestations observed in Albright hereditary osteodystrophy (AHO) and pseudohypoparathyroidism type 1A (PHP1A), including the rare occurrence of dilated cardiomyopathy observed in affected patients (*37*). Impaired PTHR signaling underlies the hallmark PTH resistance in PHP1A, leading to hypocalcemia, hyperphosphatemia, and skeletal abnormalities such as brachydactyly—features classically associated with Gαs loss-of-function mutations. Although defective V2R activation could theoretically cause subclinical vasopressin resistance, clinical signs such as nephrogenic diabetes insipidus or urine-concentrating defects are not observed in PHP1A/AHO patients with GNAS loss-of-function mutations, including those with the K53N variant (*37*, *78*). This absence of V2R-related symptoms is attributed to preserved distal tubule function (V2R-responsive), owing to biallelic Gαs expression and compensatory mechanisms such as the renin-angiotensin system. Importantly, the β2AR signaling defect associated with K53N provides a plausible molecular basis for the development of dilated cardiomyopathy: impaired Gαs-mediated β2AR cAMP production disrupts cardiac calcium handling and contractility (*73*, *79*–*81*). However, only the long isoform of Gαs K53N impairs calcium release following β-adrenergic stimulation, while paradoxically exhibiting constitutive gain-of-function cAMP production. This dual effect may contribute to cardiomyopathy through two mechanisms: (1) inhibition of calcium release, which reduces contractility and promotes pathological remodeling and reduced myocyte shortening, a hallmark of dilated cardiomyopathy (*82*–*84*), and (2) chronic cAMP/PKA overactivation, which drives maladaptive/pathological remodeling and dilated cardiomyopathy, as seen in Gαs-overexpressing mouse models with reduced contractility and dilated ventricles (*80*, *85*).

The family history and symptom segregation—particularly PHP1a manifestations (*37*)— strongly suggests that K53N mutation reside on the maternal allele of *GNAS*. PHP1a exclusively manifests with maternally inherited mutation due to genomic imprinting, as the maternal allele is preferentially expressed in key endocrine tissues including the proximal renal tubule (explaining PTH resistance), thyroid, gonads, and pituitary (*19*). Notably, this imprinting pattern extends to cardiac tissue (*30*), where dominance of maternal allele expression likely underlies the cardiomyopathy observed with K53N. Furthermore, although Gαs is ubiquitously expressed, the distribution of its short and long isoforms exhibits tissue-specific patterns across species (*16*). The long isoform predominates in the brain cortex, cerebellum, kidney, adrenal medulla, and placenta (*86*–*91*), while the short isoform is more abundant in platelets, liver, and brain striatum (*89*–*92*). Cardiac expression patterns are complex and varies by species and developmental stage: bovine heart sarcolemma favors the short isoform (*89*), whereas rabbit atrium and ventricle (adult and neonatal) (*90*) and rat neonatal cardiomyocytes (*93*) predominantly express the long isoform, consistent with our findings in NRVMs. In adult rat hearts, the atria show balanced isoform expression, while the ventricles favor the short isoform (*93*). Human data also reflects complexity: in adult right atrium, mRNA analysis indicates a predominance (∼60%) of the short isoform (*91*), but protein studies reveal strong long isoform expression (*94*). Unfortunately, human tissue/cell distribution data for Gαs isoforms remains very limited. Given the potential for isoform-specific functional differences, mutant effects, and the documented changes in splicing during development, aging, and disease (*16*, *18*), a more precise analysis of Gαs splice variant expression in human tissues is essential to fully elucidate their roles in health and disease.

In conclusion, our study reveals that the Gαs K53N variant represents a unique and complex G proteinopathy, characterized by both loss- and gain-of-function effects, dominant-negative activity, and striking isoform specificity. This duality underlies the multisystem clinical manifestations observed in patients, including classic features of PHP1a/AHO and the novel association with dilated cardiomyopathy. Our findings emphasize the critical role of alternative splicing, tissue-specific isoform expression, and genomic imprinting in modulating Gαs function and disease phenotype. Going forward, a deeper understanding of Gαs isoform distribution and regulation in human tissues will be essential for unraveling the full spectrum of GNAS-related disorders mechanisms and for developing targeted therapeutic strategies.

## Materials and Methods

### Antibodies and reagents

Anti-Gαs antibody mouse monoclonal (#sc-135914) was purchased from Santa Cruz (Dallas, Texas, USA), anti-Gαs antibody rabbit polyclonal (#371732) from Millipore-Sigma (Oakville, Ontario, Canada) and anti-GAPDH-HRP (#14C10) and -phospho-ERK (#9101) were purchased from Cell Signaling. FuGENE 6 transfection reagent was purchased from Promega (Madison, Wisconsin, USA), and Lipofectamine 3000 from Invitrogen (Waltham, Massachusetts, USA). PhosSTOP (#4906845001), cOmplete^TM^ (#11836170001), forskolin (#F3917) and cycloheximide (#239763) were purchased from Sigma. Cholera toxin (#100B) was purchased from List Labs (Campbell, California, USA), Isoproterenol (#I6504) from Sigma, oxytocin (#1910/1) was purchased from Tocris (Toronto, Ontario, Canada) and PTHrP 1-36 (#4031193) from BACHEM (Torrance, CA, USA).

### DNA constructs

pCDNA3.1-human Gαs (long and short isoforms) were obtained from the cDNA Resource Center (Bloomsburg University of Pennsylvania, PA, USA). The cAMP biosensor GFP10-EPAC-RlucII and pIRES-Gαs-RlucII(119) and pcDNA3.1-CAAX-rGFP were generous gifts from Dr. Michel Bouvier (Université de Montreal, QC, Canada). The Gαs K53N mutant (short and long isoform) were generated by replacing the lysine with asparagine at position 53 in Gαs in the plasmid pCDNA3.1 using the QuikChange site-directed mutagenesis kit (Agilent Technologies). Gαs-RlucII(119) short K53N was generated by amplifying the sequence of RlucII by PCR with the primers in table 1 from the short isoform wild type Gαs-RlucII(119) and the sequences of short isoform Gαs K53N in pcDNA3.1 vector with the primers in table 1. RlucII PCR products were next inserted into Gαs short K53N by Hifi Assembly kit (NEbuilder). pcDNA3.1-Gαs-GFP was generated based on(*95*). Primers for these different constructions are described in Table S1. All DNA constructions were sequenced before use.

### Cell culture

HEK293T (Human embryonic kidney 293) cells were kindly provided by Dr. Alexandra Newton (University of California, San Diego, CA, USA). HEK293SL is a gift from Dr. Stéphane Laporte (McGill University, QC, Canada). HEK293-Gαs-KO (HEK293 Gαs Crispr Knock-out) cell line and HEK293 parental were kindly provided by Dr. Michel Bouvier (Université de Montreal, QC, Canada) and generated as described in (*96*). All these cells were maintained in DMEM (Gibco) supplemented with 10% (v/v) FBS (Hyclone Laboratories, Logan, UT, USA) and 2 mM L-glutamine, 50 IU/mL penicillin and 50 μg/mL streptomycin (Wisent, St-Bruno, QC, Canada) at 37□ with 5% of CO_2_.

### Transfection

For Western Blot assays, HEK293-Gαs-KO were plated in 60mm dishes one day prior transfection and transfected with Lipofectamine 3000 (Invitrogen), according to the manufacturer instructions. For BRET assays, cells were seeded one day prior transfection (25×10^4^ cells per well) and transfected using linear polyethyleneimine (PEI, Polysciences, MW: 25 000) at a ratio of 3:1 (PEI:DNA ratio 3:1). For live cell confocal microscopy analysis, HEK293SL (35×10^3^ cells) were seeded in 35 mm glass-bottom dishes (MaTek Corporation) coated with Poly-L-Lysine (Sigma). Cells were transfected the day after with FuGENE 6 transfection reagent (Promega) according to the manufacturer instructions. For cAMP assays, HEK293-parental or HEK Gαs-KO cells were first seeded in a 6 well plate (20×10^4^ cells/well) and 24 hours later were transiently transfected with either no receptor (to measure response by endogenous βAR receptor), V2R (10 ng) or PTHR1 (100 ng) receptor together with Gαs or indicated Gαs mutants (150 ng) using linear polyethylenimine (PEI:DNA ratio 3:1)

### BRET assay

For cAMP EPAC biosensor experiments, cells were transiently transfected with 15ng of plasmids encoding Gαs WT or mutant Gαs, 50ng GFP10-EPAC-RlucII or receptor (10ng V2R or 30ng PTH1R). To measure the ligand induced association and dissociation of Gβγ from Gαs following GPCR stimulation, we used a short and long isoforms of Gαs, coupled to renilla luciferase at position 119 and 133, respectively (Gαs (119) S-RlucII or Gαs (133) L-RlucII), with a dimer composed of Gβ1 and Gγ1-GFP10. For this assay, cells were transiently transfected with 100ng of plasmids encoding short or long isoforms of Gαs-RlucII-WT or K53N, 500ng Gβ1, 500ng Gγ1-GFP10. A maximum of 2μg of DNA was prepared in Opti-MEM (Gibco) and Salmon sperm DNA (Invitrogen) was used to keep the same amount of total DNA. Twenty-four hours after transfection, 50 000 cells per well were distributed into white opaque 96-well plate (Falcon) and incubated for another 24 hours for BRET signal lecture. Twenty-four hours later, cells were washed with BRET buffer [146mM NaCl, 4.2mM KCl, 1mM CaCl_2_, 0.5mM MgCl2, 5.5 mM d-glucose and 10mM Hepes (pH 7.4)] and stabilized at room temperature or at 37□ (as indicated for different receptor) for 1 hour for BRET measurements. The luciferase substrate Coelenterazine 400a (GoldBio) 5μM or Prolume Purple (Nanolight technologies) 1μM was added 5min prior to measure baseline. Agonists for each receptor were added at the indicated time points respectively. All BRET signals were measured using the Infinite M1000 microplate reader (TECAN, Austria) with the energy donor filter (410 nm) for RlucII and energy acceptor filter (515 nm) for GFP10 with an integration time of 300 ms. BRET2 ratio was calculated by dividing the acceptor signal (GFP10) over the donor signal (RlucII). GraphPad Prism was used to generate kinetic curves.

### cAMP production assay

We used the HTRF cAMP Gαs Dynamic kit from Revvity Health Sciences Canada, Inc (Mississauga, Ontario, Canada) to measure cyclic adenosine monophosphate (cAMP) levels. Forty-eight hours after transfection, cells were washed with PBS at room temperature, then trypsinized and distributed at 5 000 cells/well (5 μl) in a white 384-well plate in stimulation buffer (10 mM Hepes, 1 mM CaCl_2_, 0.5 mM MgCl_2_, 4.2 mM KCl, 146 mM NaCl, 5.5 mM glucose, 0.5 mM IBMX, pH 7.4). Necessary dilutions of each ligand (2X) in addition to forskolin (10 μM) were prepared in stimulation buffer and cells were stimulated at 37°C for 30 min with indicated ligand (5 μL). Cells were then lysed with the lysis buffer containing 5 μl of cAMP coupled to the d2 dye. After addition of 5 μl of anti-cAMP cryptate terbium conjugate, cells were incubated for 3 h at room temperature under agitation. FRET signal was measured using a TECAN M1000 fluorescence plate reader (TECAN, Austria) with excitation filter (337 nm) and emission filter (620 nm). Results were normalized to the forskolin response for each condition.

### Confocal Microscopy

Cells were plated on 35 mm glass-bottom dishes (MatTek Corporation, Ashland, MA, USA) and transfected with Fugene6, as described above. 48h later, cells were starved in FluoroBrite DMEM (Gibco) for 1h at 37°C. To keep the cells at 37°C with 5% CO_2,_ Petri dishes were placed in a microchamber (Okolab, NA, Italy) attached to the stage of the microscope. 10 µM of isoproterenol (Sigma) was then added to the cells and images were taken at 0 and 10 minutes. Images were acquired with a scanning confocal microscope (Leica TCS SP8 STED DMI8, Leica Microsystems, Toronto, ON, Canada) equipped with a 63×/1.4 oil-immersion objective and a tunable white light laser (470 to 670 nm). The enhanced green fluorescent protein (EGFP) was excited with the 488 nm laser line of the white light laser and emissions were detected with a HyD detector. LAS AF Lite software (Leica) was used for image acquisition and analysis. The images were further processed using Adobe Photoshop (Adobe Systems, San Jose, CA, USA).

### Cycloheximide chase

Culture medium of transfected HEK293-Gαs-KO was removed and replaced with 2 mL of serum-free DMEM medium (Gibco) supplemented with 25 mM HEPES buffer and 40 µg/mL cycloheximide. After a 0, 0.5, 1, 2 or 4 h incubation, cells were lysed with agitation in 50 mM Tris buffer (pH 7.4) containing 150 mM NaCl, 5 mM EDTA,1% NP40, and protease inhibitors for 1h at 4°C. Cell lysates were then centrifuged 15,800x*g* for 20 min and protein concentration of the supernatants was dosed using Bradford assays (BioRad, Hercules, California, USA) and analyzed by SDS-PAGE and immunoblotting.

### Immunoblotting

Protein samples were separated on 10% SDS–polyacrylamide gel electrophoresis (PAGE) gels and transferred to 0.45 μm diameter pore-size nitrocellulose membranes (Perkin Elmer, Waltham, MA, USA). Membranes were then blocked in Tris-buffered saline (20 mM Tris-HCl, pH 7.4, 150 mM NaCl) containing 0.1% Tween 20 and 5% non-fat dry milk and were incubated with primary antibodies for 1h at room temperature and subsequently incubated with horseradish peroxidase-conjugated goat anti-rabbit or anti-mouse IgG (Bio-Rad, Hercules, CA, USA) for 1h at RT and enhanced using a chemiluminescence detection reagent (Pierce Chemical, Thermo Fisher Scientific, Waltham, MA, USA). Results are expressed as means ± SD and each experiment were performed at least in triplicate. Densitometric analyses were performed with ImageJ according to the documentation (https://imagej.nih.gov/). The statistical significance of samples was assessed using an a two-way ANOVA with a Sidak’s multiple comparisons test. A p<0.05 was considered significant.

### Neonatal rat ventricular myocytes isolation and infection

Neonatal rat ventricular myocytes (NRVMs) were extracted from the hearts of 1-3 days-old Sprague-Dawley rat pups (Charles River Laboratories) with the rat Neonatal Cardiomyocyte Isolation kit from Myltenyi Biotech (#130-105-420), as recommended by the manufacturer. Briefly, after enzymatic digestion, NRVMs were separated by a pre-plating step of 30 minutes at 37°C, where non-myocyte cells attached faster than NRVMs to the petri dish. The supernatant containing NRVMs was resuspended in M199 medium supplemented with 10% FBS, penicillin (100 U/ml)-streptomycin (100 μg/ml)-glutamine (2 mM) (PSG). For BRET assays, cells were plated on gelatin-coated 96-well white opaque tissue culture plates (Falcon, Cat No 353296) at a density of 6.6 x 10^4^ cells per well and maintained in M199 medium supplemented with 10% FBS and PSG. Twenty-four hours after plating, cells were washed twice with D-PBS containing Ca^2+^ and Mg^2+^ to remove debris before infecting NRVMs with adenoviruses encoding shRNA targeting Gαs, at a multiplicity of infection of 10, to efficiently knock-down its expression over a total period of 144h. Adenovirus were generated as previously described in (*97*). Adenovirus-mediated re-expression of wild-type or mutant forms of Gαs, at a multiplicity of infection of 5, was realized by infecting the NRVMs 2 days before performing BRET assays. The complete culture media was replaced every other day throughout the culture period in a humidified incubator with 5% CO_2_ at 37°C. For EC coupling experiments, cells were plated on fibronectin-coated 35 mm glass bottom culture dish (MatTek Corporation, Cat No P35G-1.0-20.C) at a density of 1.75 x 10^6^ cells per dish and maintained in complete media supplemented with insulin-transferrin-selenium (1x concentration, Gibco, #41400045). Twenty-four hours after plating, cells were washed twice with D-PBS containing Ca^2+^ and Mg^2+^. During the first 72h, 5-Bromo-2’-deoxyuridine (0.01mM) was added to the culture media to prevent proliferation of non-cardiomyocyte cells. Four hours after plating, cells were infected with adenoviruses encoding shRNA targeting Gαs at a multiplicity of infection of 40 over a total period of 160h. Adenovirus-mediated re-expression of wild-type or mutant forms of Gαs, at a multiplicity of infection of 5, was realized by infecting the NRVMs 3 days before performing EC coupling experiments. The complete media was replaced every 72h throughout the culture period in a humidified incubator with 5% CO2 at 37°C.

### Calcium imaging

The MultiCell High Throughput System (IonOptix) was used for the calcium imaging experiments. This system comprises a Fura-2-based dual excitation fluorescence photomultiplier system, along with a heated chamber mounted on top of a motorized microscope and an electrical stimulation assembly, which supports the measurements of rapid calcium mobilization changes from a large number of cells. Before each calcium imaging experiment, cells were serum-starved for 16 hours. Cells were washed with warm modified Tyrode buffer [NaCl 130 mM, KCl 5.4 mM, HEPES 10 mM, MgCl2 0.4 mM, CaCl2 1.8 mM, glucose 10 mM (pH 7.4)] and loaded with fura-2 AM (4μM) in buffer for 30 min at 37□. After loading, cells were washed and placed in the acquisition chamber. The temperature was maintained at 37□, and perfusion of warm buffer at a rate of 0.5 ml/min was ensured throughout the entire experiment. Electrodes inserted at the periphery of the 35 mm dish generated a 28 V impulse at a frequency of 1.7 Hz. Cells were allowed to stabilize for 3 min during the selection process of 10 cardiomyocyte patches containing 2-3 cells each displaying visible contraction. Baseline measurements were acquired for another 3 min before a bolus of warm isoproterenol solution was added to the dish to a final concentration of 1 uM. Acquisition was then resumed for an additional 3 min. The pulsed current induced by the electrode elicited contractions in the NRVMs. These contractions were associated with a transient spike in the intracellular [Ca2+] concentration, which is measurable using the integrated fluorescence system as the ratio between the fluorescence intensity of the subject when excited with 340 nm or 380 nm light. By using a 250 Hz pulsed interleaved excitation, the system measures the relative fluorescence unit (RFU) with a temporal resolution of 4 ms. The RFU traces produced by each cell during baseline and after isoproterenol stimulation were averaged and compared. For each condition, the experiment was repeated at least three times in duplicate. The response of NRVMs to a buffer bolus was only performed once.

### Calcium transient analysis

Calcium transient analysis was performed using Cytosolver 2.2.0 (IonOptix) based on the monotonic transient acquired by the IonWizard Software 7.8.3 (IonOptix). Area under the RFU transient curve values was integrated with the trapezoid method from the NumPy python library, using the averaged traces of individual cell from the Cytosolver analysis.

### Gαs protein expression and purification

The pET15b plasmid containing a human *GNAS2* gene immediately downstream of an open reading frame for hexahistidine-tagged small ubiquitin-related modifier (SUMO) and tobacco etch virus (TEV) cleavage site was kindly provided by Dr. Scott Prosser (University of Toronto, Canada). The mutation A159T (K53N) was generated via site-directed mutagenesis. The expression and purification of Gαs (identical for the WT and K53N mutant constructs) has been described previously(*98*). Briefly, vectors were transformed into *E. coli* BL21(DE3)star cells following the manufacture’s protocol. The cells were then grown in LB medium (1% tryptone, 0.5% yeast extract, 1% NaCl, supplemented with 100 μg/ml carbenicillin) until the OD600 reached 0.3. Protein expression was induced by adding 50 μM of IPTG (Sigma) and the cells were grown overnight at 19□. Cells were harvested by centrifugation at 6 000 × g for 10 minutes, and the resulting pellets were resuspended in lysis buffer containing: 50 mM Na_2_HPO_4_ (pH 8.0), 300 mM NaCl, 5 mM imidazole, 2 mM MgCl_2_, 5 mM 6-aminocaproic acid, 5 mM benzamidine, 0.4 mg/ml lysozyme, 2 μg/ml DNase I, 500 μM TCEP, 50 μM GDP and 10% glycerol. The suspension was then sonicated on ice for 5 min with 10-second interval for cooling. The lysate was clarified by centrifugation at 70 000 × g for 30 minutes. The supernatant was then loaded onto cOmplete His-tag purification resin (Roche) and washed with 20 bed volumes of wash buffer containing: 50 mM Na_2_HPO_4_ (pH 8.0), 300 mM NaCl, 25 mM imidazole, 2 mM MgCl_2_, 5 mM 6-aminocaproic acid, 5 mM benzamidine, 500 μM TCEP, 50 μM GDP and 10% glycerol. The bound protein was eluted with elution buffer containing: 50 mM Na_2_HPO_4_ (pH 8.0), 100 mM NaCl, 250 mM imidazole, 2 mM MgCl_2_, 500 μM TCEP, 50 μM GDP and 10% glycerol. The eluate was then buffer-exchanged in a buffer containing: 50 mM Na_2_HPO_4_, pH 8.0, 100 mM NaCl, 2 mM MgCl_2_ and 10% glycerol and incubated overnight at 4□ with 1.35 μM TEV protease (NEB, Ipswich, MA, USA). The cleaved product was again loaded onto cOmplete His-tag purification resin and the flowthrough was further purified by size exclusion chromatography on a Hiload 26/600 Superdex 200 prep-grade column (Cytivia) equilibrated in a buffer containing: 50 mM HEPES (pH 7.4), 100 mM NaCl, 2 mM MgCl_2_, 10% glycerol. The fractions containing folded Gαs were collected were collected and concentrated, and GDP concentration was adjusted to a molar ratio of 1.2 relative to Gαs protein. The final purified protein was snap-frozen in liquid nitrogen and stored at -80□.

### Nucleotide Exchange assay

GTP binding assay for Gαs was performed using BODIPY FL GTPγS (Invitrogen, Thermo Fisher Scientific, Waltham, MA, USA), as described in the referenced protocol (*99*), using a quartz cuvette. TEM buffer (20 mM Tris-HCl, pH 8.0; 1 mM EDTA; 10 mM MgCl_2_) containing 100 nM BODIPY FL GTPγS was used to establish the baseline fluorescence. After 100 seconds of stabilization, either control buffer, 200 nM Gαs short WT, or K53N was added, and fluorescence intensity was monitored until 2000 seconds. Measurements were performed using an F-2500 FL Spectrophotometer (Hitachi) with a 2.5 nm slit width, excitation at 488 nm, and emission at 512 nm. Data analysis and graph generation were conducted using GraphPad Prism 10.

### GTP hydrolysis assay

GTP hydrolysis assays were performed using the GTPase-Glo assay kit (Promega) following the manufacturer’s protocol. After optimizing the quantities of GTP and purified Gαs protein, 2 μM of Gαs protein and 2 μM of GTP were prepared separately in the provided buffer. The GTP hydrolysis reaction was initiated by mixing 2.5 μl of GTP with 2.5 μl of purified Gαs short WT or K53N in a 384-well AlphaPlate (PerkinElmer). A control was also carried out with only buffer and no Gαs. The reaction was terminated by adding 5 μl of the GTPase-Glo reagent, followed by incubation at room temperature for 30 minutes. This reagent converts any remaining unhydrolyzed GTP into ATP. Subsequently, 10 μl of the Detection reagent which converts ATP into a luminescent signal was added, and the mixture was incubated for an additional 5–10 minutes at room temperature. The resulting luminescence (which is proportional to the amount of unreacted GTP) was measured using an Infinite M1000 (TECAN) plate reader with an integration time of 1000 ms. Data analysis and graph generation were conducted using GraphPad Prism 10. The amount of GTP hydrolysis for each condition (x) can be represented by its difference in luminescence with the control: % GTP hydrolysis= (Lum_0_ – Lum_x_) x100, where Lum_0_ is the luminescence signal intensity of the control reaction.

### Statistical analysis

Statistical analysis was performed using Prism version 8.2.1 or 9.3.0 (GraphPad Software) using a mixed-effects analysis (restricted Maximum Likelihood, REML) was performed with a Tukey correction for multiple comparisons (figure 2C). A one-way ANOVA multiple comparisons with Šídák correction was performed to study Gαs and Gβ1γ1 interaction (Figure 7C). For cAMP quantification assay (Figure 3C, 4B and 5B), a two-way ANOVA multiple comparisons with Šídák correction was performed. For calcium mobilization experiments (Figure 9B), a matched measure one-way ANOVA multiple comparison with Tukey correction was performed. Data were considered significant when *p* values were <0.05.

## Acknowledgments

We are grateful to Stephane Laporte (McGill University, Canada) for providing the HEK293SL cell line and to Asaka Inoue (Tohoku University, Japan) for providing the HEK293T-Gαs KO cell line. We thank Michel Bouvier (IRIC, Canada) for sharing various BRET biosensors. We thank Véronique Blais (Universtié de Sherbrooke) for her help to set up the purification of Gαs protein. We would like to acknowledge the Plateforme de microscopie photonique de l’Université de Sherbrooke for their support.

## Funding

This research was funded by the Canadian Institutes of Health Research (CIHR) to CL, MAM, RL and J-LP (grant # 191933). Additionally, AL received predoctoral fellowships from the FRQNT-funded PROTEO Network and the Fonds de Recherche du Québec (FRQS). J.M. received the Louis Gendron and Mylène Côté Excellence Scholarship from Fondation de l’Université de Sherbrooke (FUS) and a Canada Graduate Scholarships – Master’s (CGS M) award from the Canadian Institutes of Health Research (CIHR). M.A.M. was supported by a FRQS–Senior research scholarship-career award (313286).

## Author Contributions

H.G. planned and performed most of the experiments, collected and analyzed the data, prepared most of the figures, and drafted the manuscript. B.H. carried out the CAAX-BRET assays and HT-FRET cAMP assays and contributed to manuscript revision. J.M. isolated neonatal rat ventricular myocytes (NRVMs) and performed calcium-imaging experiments and transient-analysis. H.G. also contributed to NRVM isolation. E.L performed the cycloheximide assay. A.L. performed live-cell confocal imaging assays. A.D. help with the purification of Gαs. R.L. and J.L.P. provided intellectual input and revised the manuscript. L-P.P. guided the purification of Gαs and the biochemical assays, generated the Gαs structural figure, contributed to data interpretation, and revised the manuscript. M.A.M. help designed the NRVM studies, contributed to data interpretation, and revised the manuscript. C.L. conceived the overall study, provided intellectual guidance, contributed to data interpretation and wrote the manuscript. All authors approved the final version.

## Competing interests

The authors declare that they have no competing interests.

## Data and materials availability

All data needed to evaluate the conclusions in the paper are present in the paper and/or the Supplementary Materials.

---

**Table S1.**
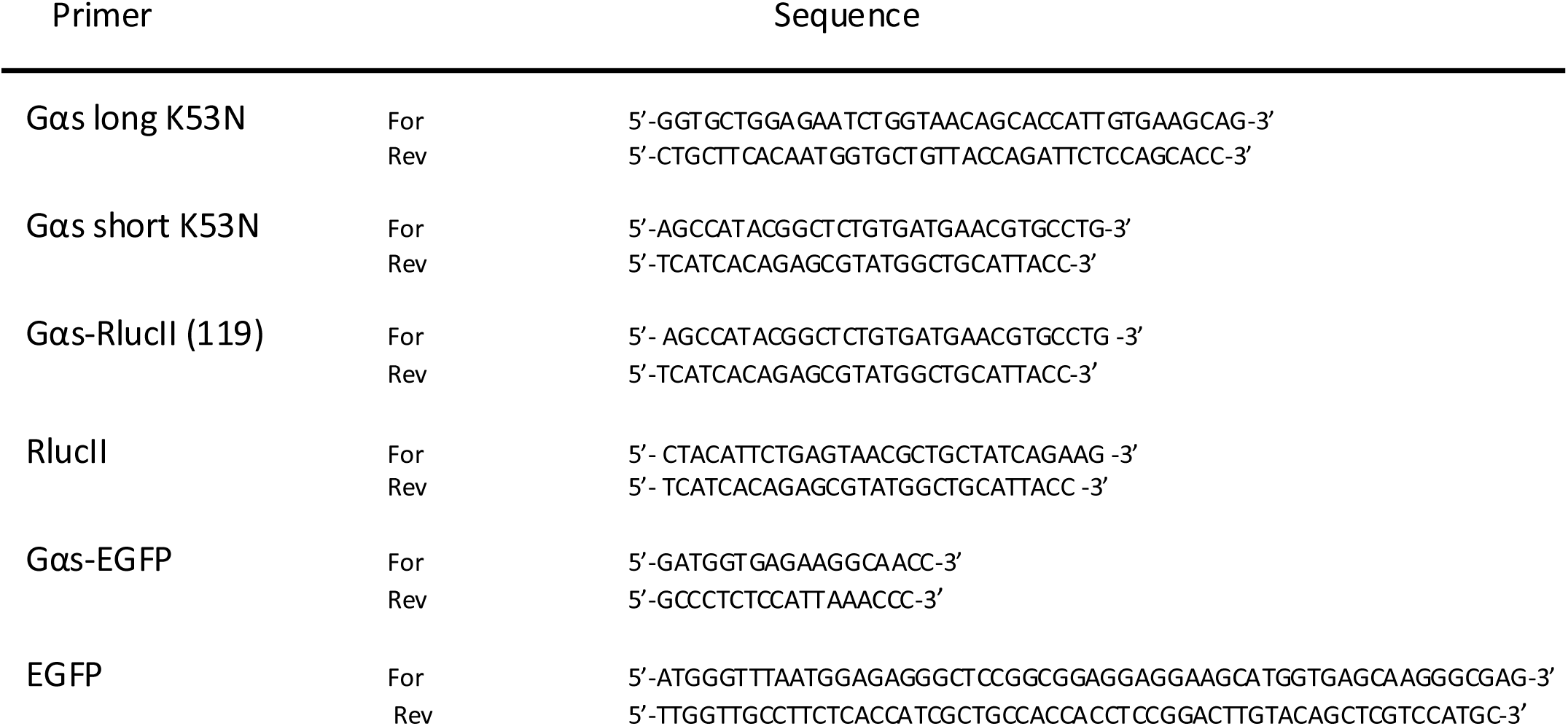
Primers used to generate the construction described in section material and method.

**Figure S1.**
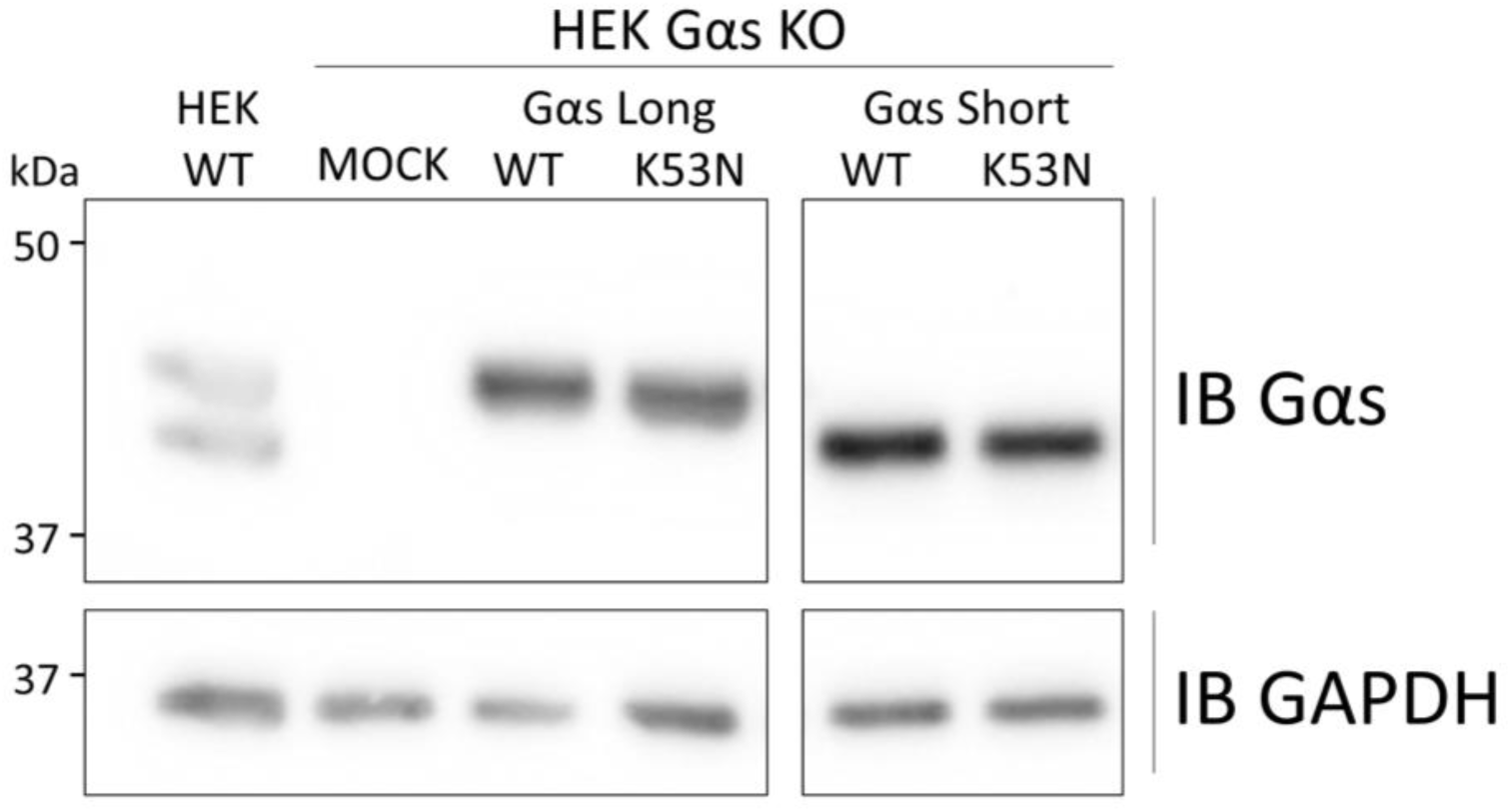
Similar expression level of Gαs in lysates from HTFR assay samples. Representative Western blot of cell lysate of Figure 3B. All samples were run on the same gel and blotted; the boxes indicate that intervening lanes were digitally removed to juxtapose samples that were originally separated on the membrane.

**Figure S2.**
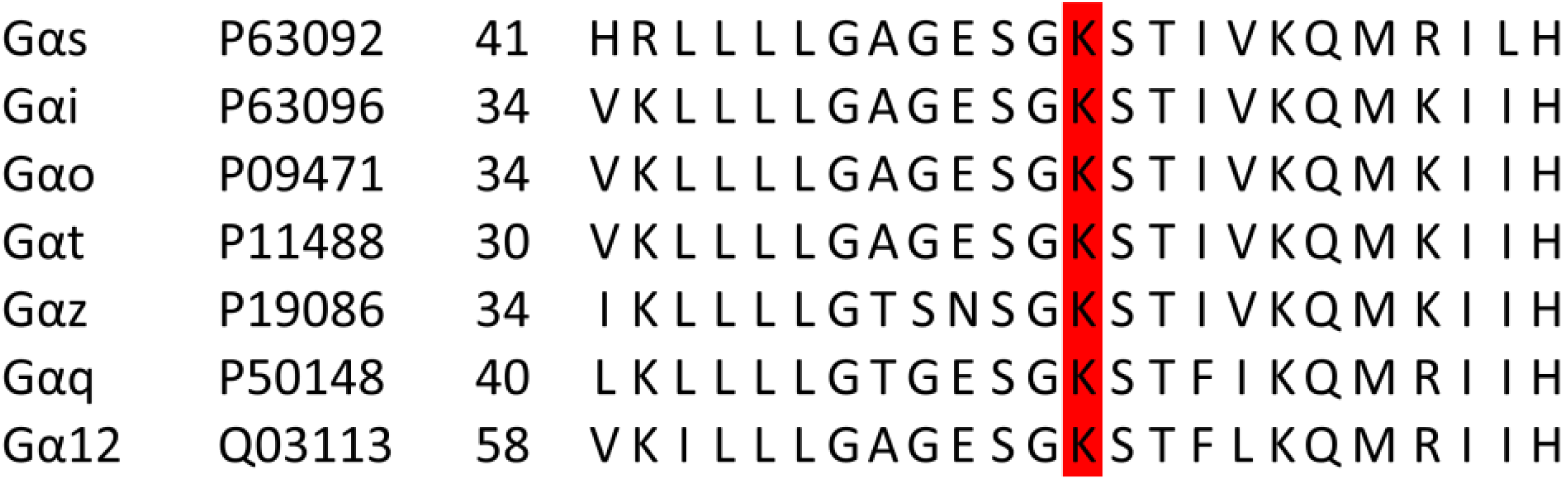
Alignment of the 41-64 region of Homo Sapiens Gαs long (p63092) with other Gα protein indicating the highly conserved region and K53 residue.

## Notes

### Competing Interest Statement

The authors have declared no competing interest.

