## Supplementary Figure S1, S2 and Table S1 for "Isoform-Dependent Loss- and Gain-of-Function of the Gαs K53N Variant in Human Disease"

Haoran Geng *et al.*

**This PDF file includes:**

Figures: S1 to S2

Tables: S1

Fig. S1.

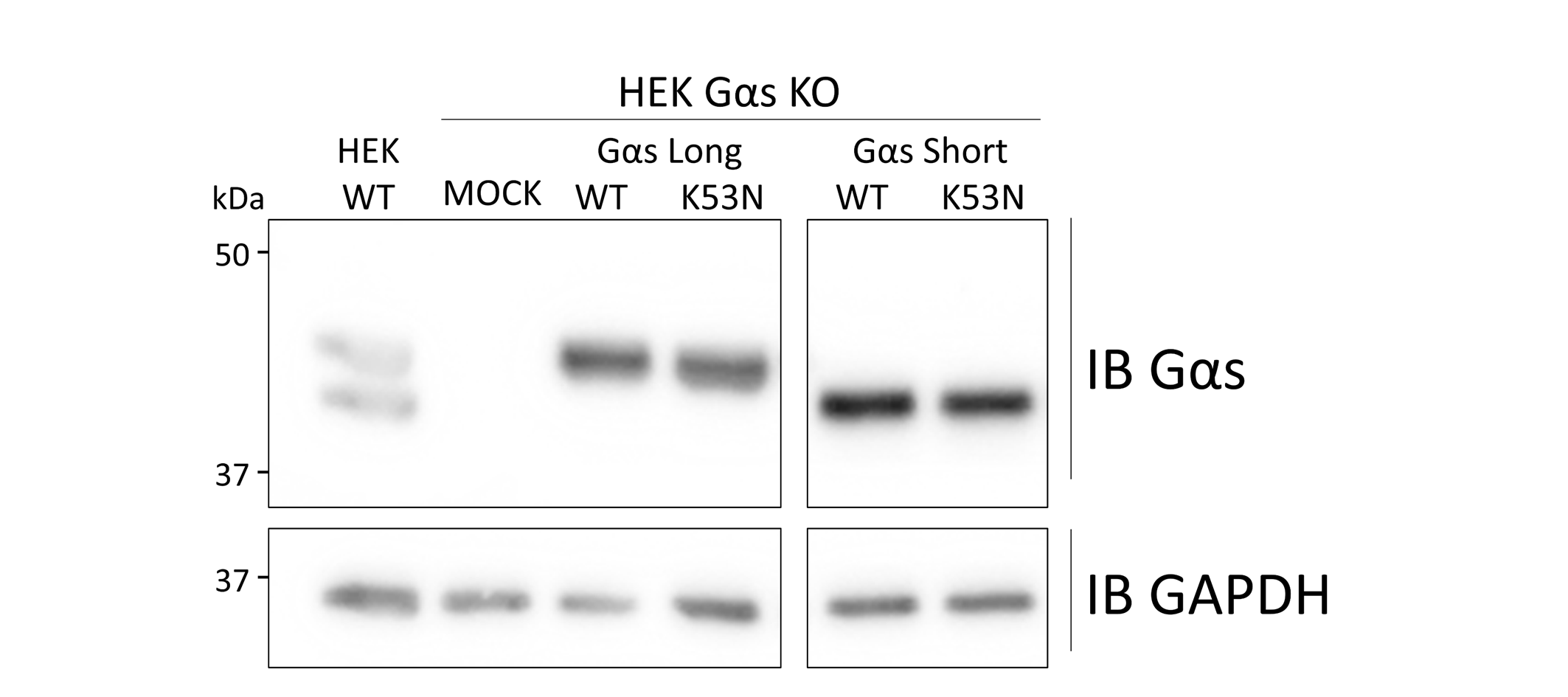

**Similar expression level of Gαs in lysates from HTFR assay samples.** Representative Western blot of cell lysate of Figure 3B. All samples were run on the same gel and blotted; the boxes indicate that intervening lanes were digitally removed to juxtapose samples that were originally separated on the membrane.

Fig. S2.

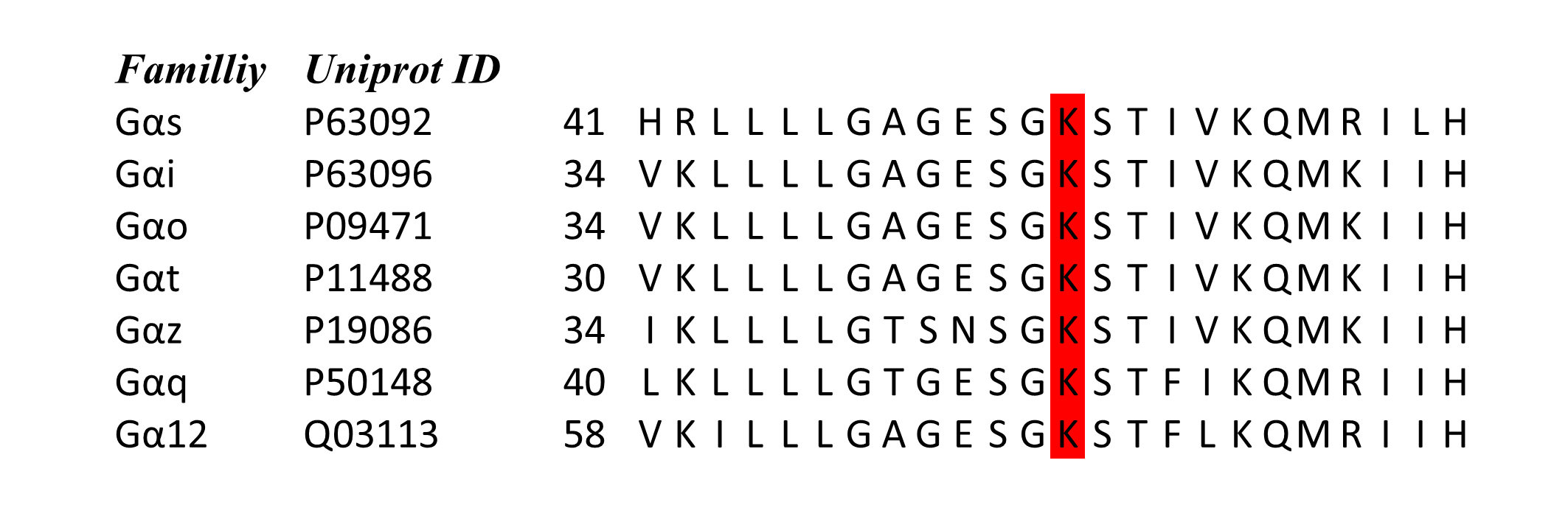

**Alignment of the 41-64 region of Homo Sapiens Gαs long (p63092) with other Gα protein indicating the highly conserved region and K53 residue.**

Table S1.

| Primer |  |  | Sequence |
| --- | --- | --- | --- |
| Gαs long K53N |  | For | 5’-GGTGCTGGAGAATCTGGTAACAGCACCATTGTGAAGCAG-3’ |
|  |  | Rev | 5’-CTGCTTCACAATGGTGCTGTTACCAGATTCTCCAGCACC-3’ |
| Gαs short K53N |  | For | 5’-AGCCATACGGCTCTGTGATGAACGTGCCTG-3’ |
|  |  | Rev | 5’-TCATCACAGAGCGTATGGCTGCATTACC-3’ |
| Gαs-RlucII (119) |  | For | 5’- AGCCATACGGCTCTGTGATGAACGTGCCTG -3’ |
|  |  | Rev | 5’-TCATCACAGAGCGTATGGCTGCATTACC-3’ |
| RlucII |  | For | 5’- CTACATTCTGAGTAACGCTGCTATCAGAAG -3’ |
|  |  | Rev | 5’- TCATCACAGAGCGTATGGCTGCATTACC -3’ |
| Gαs-EGFP |  | For | 5’-GATGGTGAGAAGGCAACC-3’ |
|  |  | Rev | 5’-GCCCTCTCCATTAAACCC-3’ |
| EGFP |  | For | 5’-ATGGGTTTAATGGAGAGGGCTCCGGCGGAGGAGGAAGCATGGTGAGCAAGGGCGAG-3’ |
|  |  | Rev | 5’-TTGGTTGCCTTCTCACCATCGCTGCCACCACCTCCGGACTTGTACAGCTCGTCCATGC-3’ |

**Primers used to generate the construction described in section material and method.**
